# The evolution of tumour composition during fractionated radiotherapy: implications for outcome

**DOI:** 10.1101/175828

**Authors:** Thomas D. Lewin, Philip K Maini, Eduardo G Moros, Heiko Enderling, Helen M Byrne

**Affiliations:** Wolfson Centre for Mathematical Biology, Mathematical Institute, University of Oxford, UK; Radiation Oncology, H. Lee Moffitt Cancer Center & Research Institute, USA; Integrated Mathematical Oncology, H. Lee Moffitt Cancer Center & Research Institute, USA

## Abstract

Current protocols for delivering radiotherapy are based primarily on tumour stage and nodal and metastases status, even though it is well known that tumours and their microenvironments are highly heterogeneous. It is well established that the local oxygen tension plays an important role in radiation-induced cell death, with hypoxic tumour regions responding poorly to irradiation. Therefore, to improve radiation response, it is important to understand more fully the spatiotemporal distribution of oxygen within a growing tumour before and during fractionated radiation. To this end, we have extended a spatially-resolved mathematical model of tumour growth first proposed by Greenspan (Stud. Appl. Math., 1972) to investigate the effects of oxygen heterogeneity on radiation-induced cell death. In more detail, cell death due to radiation at each location in the tumour, as determined by the well-known linear-quadratic model, is assumed also to depend on the local oxygen concentration. The oxygen concentration is governed by a reaction-diffusion equation that is coupled to an integro-differential equation that determines the size of the assumed spherically-symmetric tumour. We combine numerical and analytical techniques to investigate radiation response of tumours with different intra-tumoral oxygen distribution profiles. Model simulations reveal a rapid transient increase in hypoxia upon re-growth of the tumour spheroid post-irradiation. We investigate the response to different radiation fractionation schedules and identify a tumour-specific relationship between inter-fraction time and dose per fraction to achieve cure. The rich dynamics exhibited by the model suggest that spatial heterogeneity may be important for predicting tumour response to radiotherapy for clinical applications.

## 1 Introduction

Cancer continues to be one of the main causes of mortality worldwide, with 8.2 million cancer-related deaths estimated to have occurred globally in 2012 (Torre et al, 2015). Of the millions of people diagnosed with some form of cancer each year, about half will receive radiotherapy as part of their treatment (Fowler, 2006).

Typically, total radiation dose is delivered to the tumour as a series of small doses, or fractions, administered over a period of several days or weeks in order to limit the toxic side effects to healthy cell populations. The conventional fractionation schedule for most tumors, where a dose of approximately 2 Gy (Gray) is delivered once a day Monday to Friday up to a total of 50-70 Gy, has remained the standard of care for many years (Ahmed et al, 2014; Marcu, 2010). Whilst altered schemes, such as hyperfractionation, accelerated fractionation and hypofractionation, have been suggested as alternatives for certain indications, the selection of an optimal fractionation protocol for a particular tumour would clearly benefit from a more individualised approach. In particular, beyond tumour location and stage, patient-specific factors that may be important in determining response to a particular treatment are currently not considered. It is in this regard that mathematical modelling has the potential to play an important role; identifying key factors that determine treatment outcomes and ultimately identifying patient-specific treatment protocols.

Radiation damages to a DNA fragment can be either a single- or double-strand break (Hendry, 1979; Thrall, 1997; Muriel, 2006). A normal cell has repair mechanisms designed to correct for single-strand breaks. However these can fail, with misrepair potentially leading to cell death. Whilst double-strand breaks are rarer than single-strand breaks, they are more likely to result in cell death due to increased difficulty in repairing the damaged DNA. Tumours differ in their responses to irradiation, with radio-sensitivity being understood as an intrinsic property of the cell population that could be estimated from molecular analysis of biopsy samples (Eschrich et al, 2009). Other factors that affect a tissue’s response to radiotherapy include repopulation between radiation doses, tissue re-oxygenation and redistribution of cells within the cell cycle (Thrall, 1997). For rapidly proliferating tumours in particular, repopulation between fractions of radiation is important. For tumours containing larger regions of hypoxia, re-oxygenation of the tumour may play a pivotal role. Ionising radiation interacts with oxygen and water molecules, creating oxygen free radicals that are then able to cause DNA damage (indirect action). Thus, tumour cells in hypoxic regions exhibit decreased response to radiotherapy.

Due to the difficulties of studying tumours *in vivo*, such effects are often investigated experimentally using *in vitro* multicellular tumour spheroids. Tumour spheroids provide a controlled study environment of intermediate complexity between 2D culture media and *in vivo* models in which important insight can be gained into the biology of the tumour micro-environment and the effect of therapies on a growing tumour (Carlson et al, 2006; Hirschhaeuser et al, 2010; Mueller-Klieser, 1987; Sutherland et al, 1981). However, tumour spheroid growth is avascular with a typical diameter of less than 5*mm*, so while useful for calibrating mathematical models, parameters derived from tumour spheroids may need adjustment before they can be used to simulate *in vivo* tumours.

Clonogenic assays are widely used to determine clonogenic survival for a particular cell type after acute radiation doses (Joiner and van der Kogel (2009), Chapter 4). Measurements of the clonogenic surviving fraction for a single cell type across a range of doses can be used to generate a survival curve. Such curves are of similar shape across most cell types and give rise to the most widely adopted mathematical model of radiotherapy response, the linear-quadratic (LQ) model ((Joiner and van der Kogel, 2009), Chapter 4). We note that although this model is empirical, mechanistic models have been proposed to explain it ((Sachs et al, 1997; Nilsson et al, 1990), (Joiner and van der Kogel, 2009) Chapter 4). In the LQ model, the survival fraction, *SF_d_*, of tumour cells after a dose, *d* Gy, of radiationis given by

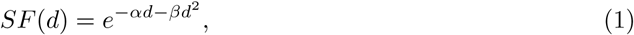

where *α* (Gy^-1^) and *β*(Gy^-2^) are intrinsic radio-sensitivity parameters (Fowler, 2006; Sachs et al, 1997; Withers, 1999; O’Rourke et al, 2009). Values for *α* and *β* may differ significantly between tumours and patients. The *α/β* (Gy) ratio is often used to characterize the sensitivity of a particular tissue type to fractionation. Values of *α/β* can fall as low as 1 Gy for late-responding tissues such as prostate cancer, and reach as high as 20 Gy for early-responding, rapidly proliferating tissues such as head and neck cancer (Withers, 1999). The LQ model is frequently incorporated into more detailed tumour models as an instantaneous effect.

Many different mathematical models and methodologies have been used to study tumour growth. Among the simplest of these are ordinary differential equation (ODE) models that aim to qualitatively capture observed growths. Small, initial tumour volumes often exhibit exponential growth dynamics, followed by a deceleration as tumour growth saturates due to microenvironmental effects, such as limited space and nutrient supply, resulting in sigmoidal growth curves (Sachs et al, 2001; McAneney and O’Rourke, 2007). Popular ODE models include the logistic and Gompertz growth models where the volume-saturation limit is represented by the carrying capacity, *K*.

Mathematical models are often built on the simplifying assumption that a tumour has a spatially homogeneous composition. Phenomenological models contain few parameters and do not account for complex underlying biological interactions. Whilst such models may provide limited biological insight, the ease with which model parameters may be estimated from limited clinical data makes them attractive for making predictions about radiotherapy response and identifying personalised fractionation protocols in the clinic (Prokopiou et al, 2015).

A variety of more complex, spatially-resolved, models have been proposed in order to provide more mechanistic insight, often reflecting the heterogeneous nature of growing tumours. As a tumour grows, it will typically develop hypoxia and necrosis in regions where oxygen or nutrient supply is inadequate. Multicellular, avascular tumour spheroids are used as *in vitro* models to study, in a controlled environment, effects observed in *in vivo* tumours (Carlson et al, 2006; Hirschhaeuser et al, 2010; Mueller-Klieser, 1987; Sutherland et al, 1981). In a similar manner, it is natural to develop mathematical models for avascular tumour spheroids before considering the more complex case of vascular tumour growth.

One of the simplest spatially-resolved models of avascular tumour growth was proposed by Greenspan (Greenspan, 1972). In the Greenspan model, the outer tumour radius evolves in response to a single diffusible nutrient species, commonly taken to be oxygen. Internal free boundaries, decomposing the tumour spheroid into a central necrotic core, an outer proliferating rim and an intermediate hypoxic annulus, are determined by contours of the oxygen profile across the tumour.

Under certain simplifying assumptions, Greenspan’s model can be reduced to a coupled system of ODEs and algebraic equations. Most other spatially-resolved continuum models are formulated as systems of partial differential equations (PDEs). Such approaches include frameworks from multiphase modelling and morphoelasticity. Multiphase models consider the tumour environment as a mixture of two or more separate fluid cell populations, or phases. Systems of PDEs are obtained by applying mass and momentum balances to each phase and making suitable constitutive assumptions about their properties and interactions (Breward et al, 2003; Byrne et al, 2003; Preziosi and Tosin, 2009). Morphoelastic models of tumour growth provide a theoretical framework within which to study biological tissues for which growth and elasticity are inter-related (Araujo and McElwain, 2004).

While incorporating more biological detail, these more sophisticated approaches typically involve more parameters that may be difficult to estimate in practice. In this paper we present a simple spatially-resolved model for tumour spheroid growth and response to radiotherapy in order to investigate the effects of spatial heterogeneity. In Section 2 we extend Greenspan’s original spatial model for tumour spheroid growth (Greenspan, 1972) to include radiation effects. In Section 3.1 we numerically solve the model equations and discuss key features of the resulting dynamics. We use a combination of further numerical simulations and analysis of the model equations in order to explore the tumour dynamics exhibited by the model.

## 2 Model development

In this section, we introduce a spatial model of avascular tumour growth originally developed by Greenspan (Greenspan, 1972). We extend Greenspan’s model to account for the effects of radiation and arrive at a new model for tumour response to radiotherapy.

### 2.1 Original growth model by Greenspan

Following Greenspan (Greenspan, 1972), we consider the growth of a radially-symmetric, avascular tumour spheroid in response to a single, growth-rate limiting, diffusible nutrient, here oxygen. For simplicity, we assume that growth inhibition occurs due to nutrient deficiency rather than in response to an inhibitory factor. We denote the outer tumour radius by *R*(*t*) and let *c*(*r, t*) denote the oxygen concentration at a distance 0 ≤ *r* ≤ *R*(*t*) from the tumour centre. We suppose that the oxygen concentration is maintained at a constant level, *c* _*∞*_, on *r* = *R*(*t*). Oxygen diffuses on a much shorter timescale than tumour growth,taking ∼10 seconds to diffuse 100 *μm* by comparison with a tumour growth period measured in days (Greenspan, 1972). As such we assume that the oxygen concentration is in a quasi-steady state. Internal free boundaries at *r* = *R*_*H*_ (*t*) and *r* = *R_N_* (*t*) are defined implicitly and denote contours on which the oxygen concentration attains the threshold values *c_H_* and *c_N_*, respectively. These interfaces decompose a well-developed tumour into a central necrotic core (0 < *r* < *R_N_*) where *c* ≤ *c_N_*, and an outer, oxygen-rich region (*R*_*H*_ < *r* < *R*) in which *c_H_* < *c*, these regions being separated by a hypoxic annulus (*R_N_* < *r* < *R*_*H*_) in which *c_N_* < *c* < *c_H_* (see Figure 1 for a schematic).

**Figure 1:**
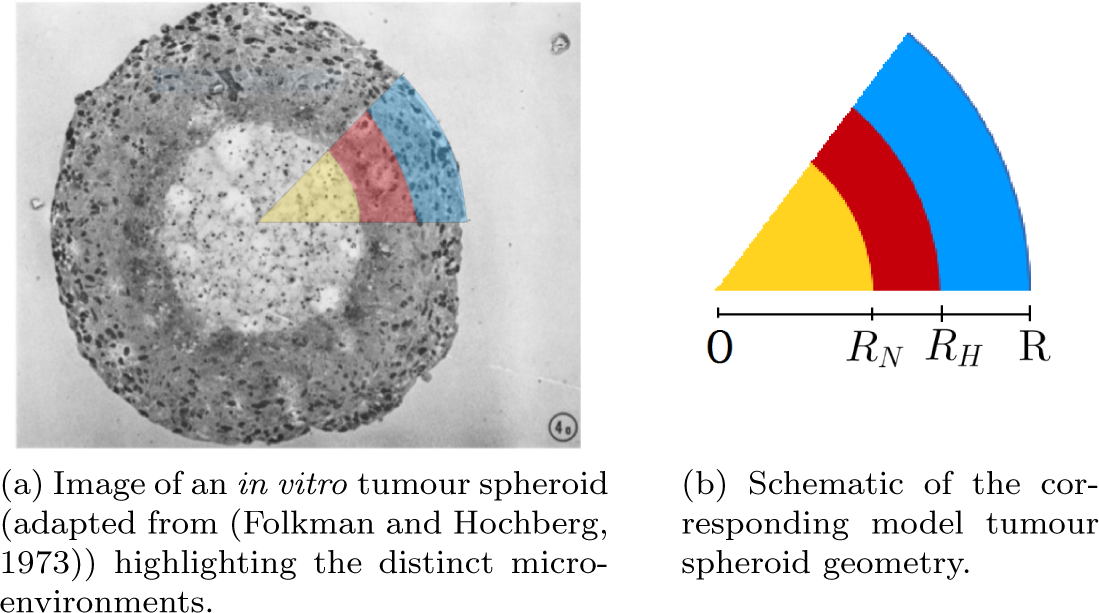
Diagrams highlighting the distinct regions within a tumour spheroid and how they are influenced by the oxygen profile. A central necrotic core (0 < *r* < *R_N_*) is surrounded by a hypoxic annulus (*R_N_* < *r* < *R*_*H*_) and an outer proliferating rim (*R*_*H*_ < *r* < *R*). The moving boundaries *r* = *R_N_* (*t*), *R*_*H*_ (*t*) and *R*(*t*) delineate these regions. Greenspan’s original model (Greenspan, 1972) describes how the dependent variables *c*(*r, t*), *R*(*t*), *R*_*H*_ (*t*) and *R_N_* (*t*) evolve over time and can be written in dimensionless form as follows:

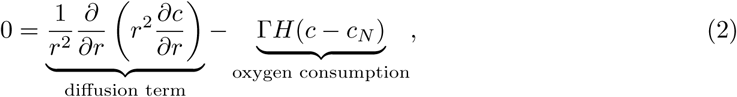

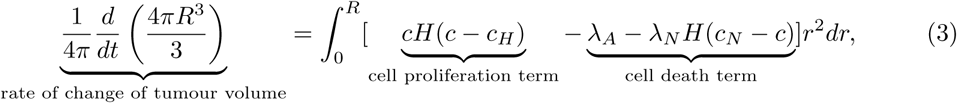

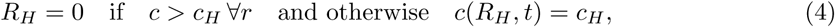

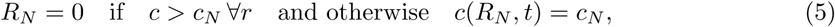

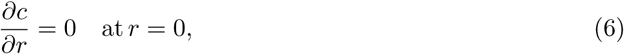

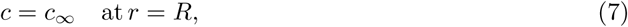

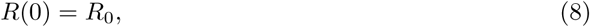

Where *H*(.) is the Heaviside function (*H*(*x*) = 1 for *x* >0, and *H*(*x*) = 0 otherwise), and Γ *λ_A_*, *λ_N_*, *c*_∞_, *c*_*H*_ & *c*_*N*_ are positive constants. Equations (2)-(8) may be reduced to a system involving a single ODE for *R*(*t*) and algebraic equations for *R*_*H*_ (*t*) and *R*_*N*_ (*t*). Analysis of the resulting equations and details of the non-dimensionalisation may be found in (Greenspan, 1972; Byrne, 2012), and for further explanation of the underlying model assumptions see Appendix A.1. Figure 2 shows how the size and composition of the tumour spheroid evolve for a typical simulation of the Greenspan model using the dimensionless parameter values in Table 1. A table of dimensional parameter values is given in Appendix A.1. Where possible we have obtained parameter estimates for model simulations from the literature, however we note that some of these values pertain to estimates used in other modelling scenarios. In this paper though, specific parameter combinations serve only to highlight the dynamics of the model and, in particular, the resulting model analysis is independent of the choice of parameter values. The effects of some of the parameters on the growth dynamics and steady state tumour composition are explored in Appendix A.2.

**Table 1:** Dimensionless parameter values for Greenspan’s model of tumour spheroid growth.

| Parameter | Symbol | Value | Source |
| --- | --- | --- | --- |
| Oxygen consumption rate | $\Gamma$ | 1.1051 | Grimes et al (2014) |
| Oxygen concentration at tumour boundary | $c_\infty$ | 1 | Grimes et al (2014) |
| Hypoxic oxygen threshold | $c_H$ | 0.1 | Grimes et al (2014) |
| Anoxic oxygen threshold | $c_N$ | 0.008 | Grimes et al (2014) |
| Apoptosis constant | $\lambda_A$ | 0.32 | Frieboes et al (2007) |
| Necrosis constant | $\lambda_N$ | 0.0061 | Schaller and Meyer-Hermann (2006) |

**Figure 2:**
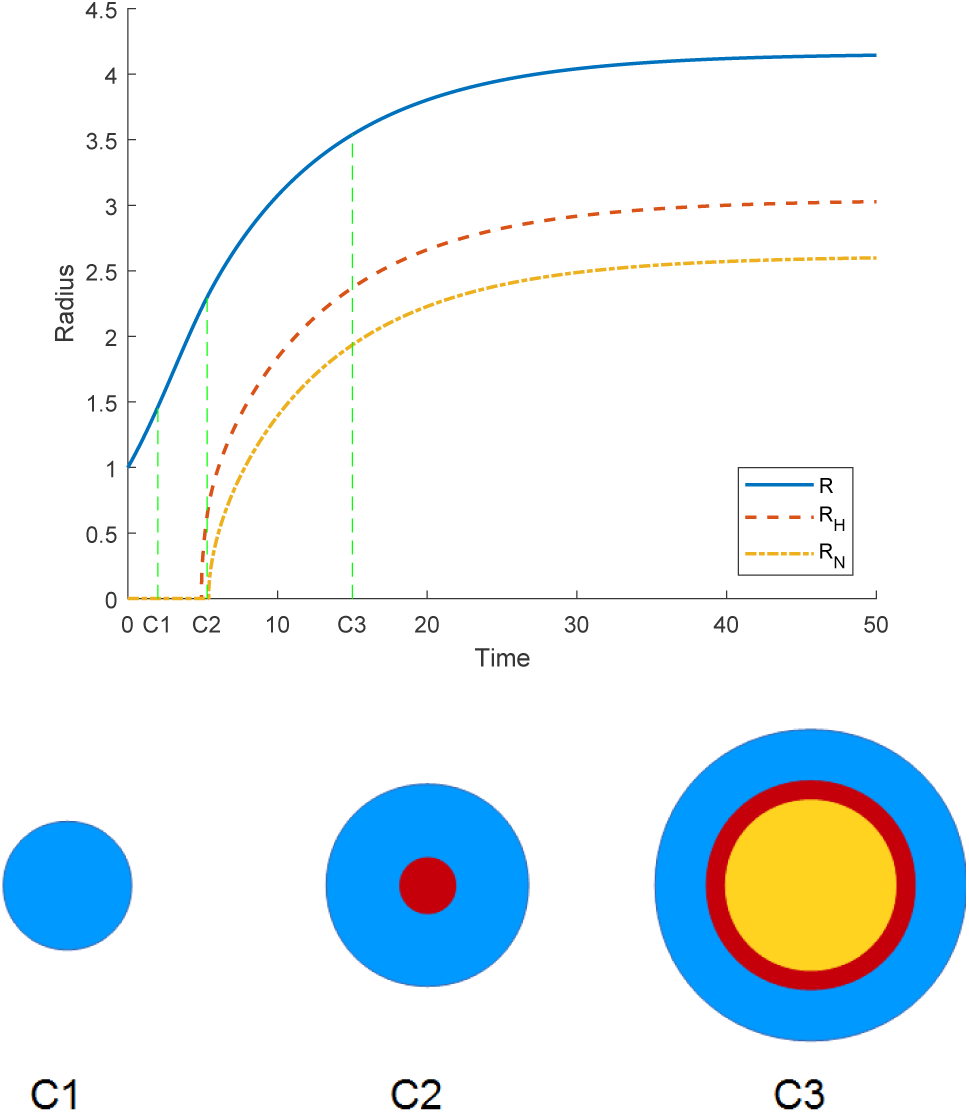
Tumour spheroid composition at various stages of tumour growth under the Greenspan model (Equations (2)-(8)) using the parameter values in Table 3; blue solid line proliferating cells, red dashed line hypoxic region, yellow dot/dashed line necrotic core

### 2.2 Incorporating radiotherapy into the Greenspan model

We adapt Greenspan’s model to account for radiotherapy by assuming that when a dose of radiotherapy is applied it causes an instantaneous dose-dependent death of viable cancer cells and, thus, a change in the size and composition of the tumour. The efficacy of the radiation and subsequent cell death depends on the local oxygen concentration. As such, we determine the radiation-induced cell death in each tumour region separately. We denote by *R*^±^, 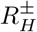and 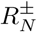the radii immediately before (-) and after (+) radiotherapy in the proliferating, hypoxic and necrotic tumor compartments.

#### 2.2.1 Radiation-induced cell death and the Linear-Quadratic model

We use the linear-quadratic model (Fowler, 2006; Joiner and van der Kogel, 2009; Enderling et al, 2010) to account for cell kill due to radiotherapy in the well-oxygenated outer rim, *R*_*H*_ < *r* < *R*, so that the volume survival fraction, *SF* (*d*), immediately after a dose *d* of radiation is given by

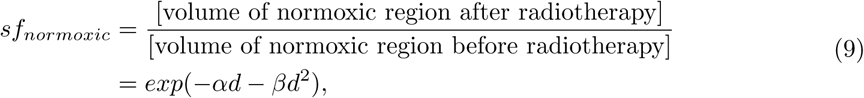

for radio-sensitivity parameters *α* and *β*. Values of *α/ β* vary markedly, with typical values falling in the range of 3-10 Gy. However extremal values of 1 Gy and 20 Gy have been reported for late-responding tissues such as prostate cancer and early-responding, rapidly proliferating tissues such as head and neck cancer, respectively (Withers, 1999). Typical radio-sensitivity parameters for rapidly-proliferating, early-responding tumours are shown in Table 2.

**Table 2:** Typical radio-sensitivity parameters

| Parameter | Value | Source |
| --- | --- | --- |
| $\alpha$ (Gy<sup>-1</sup>) | 0.35 | Fowler (2006); Muriel (2006); Sachs et al (2001) |
| $\alpha/\beta$ (Gy) | 10 | Withers (1999) |
| OER (dimensionless) | 3 | Carlson et al (2006) |

When exposed to ionising radiation, the potent oxygen free radicals that form in well-oxygenated regions increase the amount of DNA damage by up to a factor of 3 when compared with hypoxic tumour regions (Alper and Howard-Flanders, 1956). Following Carlson *et al.* (Carlson et al, 2006), we incorporate the oxygen enhancement ratio, OER, to account for this effect. Radio-Sensitivity parameters are typically quoted for normoxic conditions. As such we take the common approach and use the OER as a constant factor (taken to be equal to 3 in the presented simulations) that reduces the intrinsic radio-sensitivity parameters of tumour cells, *α* and *β*, in hypoxic regions. This creates a discontinuity in the response to radiotherapy at *r* = *R*_*H*_. The volume survival fraction immediately after a dose of radiation is delivered to a population of hypoxic cells is given by

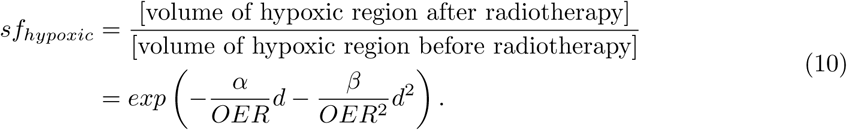

There is no clear consensus in the literature as to how to modify the ‘quadratic’ component of cell death, however the form used in Equation (10) affords the interpretation of the OER as the multiplying factor for the dose escalation required under hypoxia in order to achieve the same cell kill as under normoxic conditions. We note that, in practice, this effect is likely to depend continuously on the local oxygen concentration. A continuous functional form for the OER (*OER* = *OER*(*c*)) was proposed by Alper & Howard-Flanders (Alper and Howard-Flanders, 1956). For realistic parameter regimes, simulations using the continuous OER presented in (Alper and Howard-Flanders, 1956) and the discrete OER presented here did not significantly vary (results not shown). For these reasons we restrict attention to the discrete OER stated above.

We assume further that dead material within the necrotic core is unaffected by radiation and so we impose that 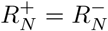.

Combining these assumptions we deduce that, for a well-developed tumour spheroid (Figure 2, case C) with 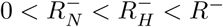, a dose *d* of radiation gives a total volume change of

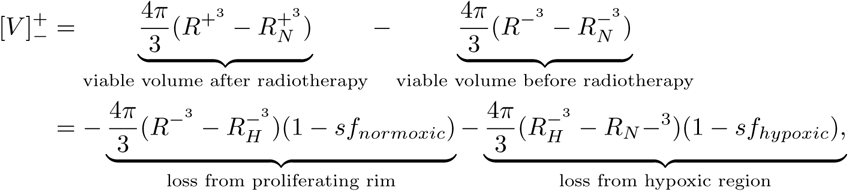

which, on rearrangement, yields the following expression for *R*^+^ in terms of *R*^-^, 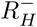and 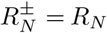:

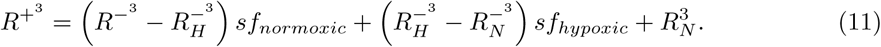

The oxygen concentration profile associated with the new tumour structure can be determined (Equation (2)) and *R*_*H*_ defined implicitly as before (see Equation (4)). We note that while not all hypoxic tumour cells will be killed from a single radiotherapy fraction (*sf*_*hypoxic*_ > 0), the instantaneous volume loss due to irradiation and the subsequent reoxygenation of the tumour spheroid may result in a post-radiotherapy tumour composition without a hypoxic region. Two cases can arise when a well-developed, 3-layer tumour is irradiated (see Figure 3):

(i) 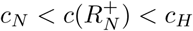 - irradiated tumour is a fully-developed tumour spheroid with 3 layers;

(ii)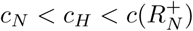 - irradiated tumour has a necrotic core, but no hypoxic annulus.

Since the necrotic core is assumed to be unaffected by radiotherapy, these are the only possible options. We have already described how a tumour spheroid of case (i) responds to radiation (Equation (11)). If a tumour with a pre-radiotherapy composition as in case (ii) is irradiated then the corresponding volume change is given by

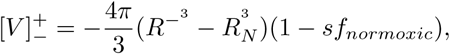

which, on rearrangement, supplies the following expression for *R*^+^ in terms of *R*^-^ and *R*_*N*_

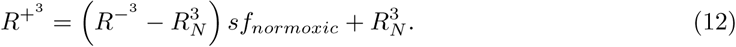

**Figure 3:**
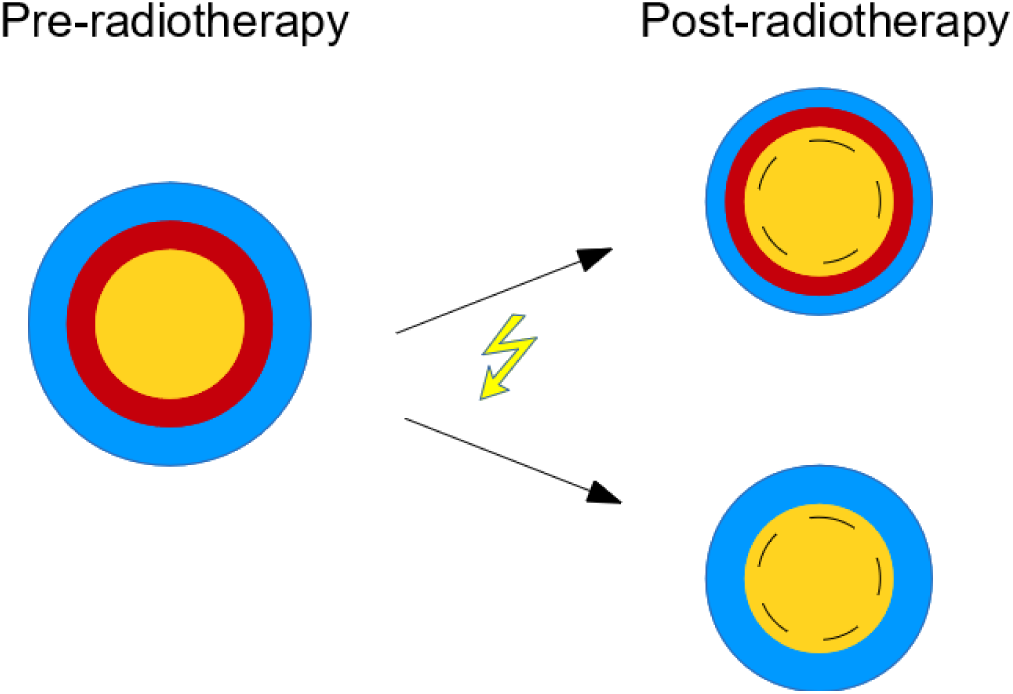
Schematic of the model tumour spheroid post-radiotherapy. Dashed line represents new position of the internal contour *c* = *c_N_* and shows the resulting mismatch between this contour and *R_N_*.

#### 2.2.2 Reconciling preand post-radiation tumour growth

In the normal growth regime without radiotherapy, *R*_*N*_ is defined implicitly by the oxygen concentration (see Equation (5)). When the tumour volume is reduced due to radiation, the oxygen concentration at the centre of the tumour increases and the location of the interface on which *c* = *c*_*N*_ will shift towards the tumour centre or disappear. As the necrotic core is unaffected by radiation, 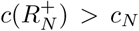and the growth of the tumour spheroid immediately post-radiotherapy does not follow the original, control dynamics.

Between fractions of radiotherapy, repopulation of the tumour occurs. The tumour cells proliferate and die as before, however, whilst *c*(*R*_*N*_) > *c*_*N*_ no new material is added to the necrotic core, and so the existing necrotic material simply decays at the rate *λ_A_* + *λ_N_*. In this case *R*_*N*_ evolves according to

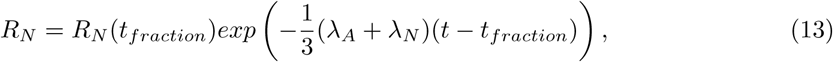

where *t*_*fraction*_ is the time of the last fraction delivered for which *c*(*R*_*N*_) = *c*_*N*_, and *R*_*N*_ (*t*_*fraction*_) is the radius of the necrotic core upon irradiation. As such, prior to treatment with radiotherapy at *t* = *t_fraction_* the tumour composition is consistent with the control, Greenspan dynamics, while Equation (13) governs the evolution of *R*_*N*_ post-irradiation.

Equations (2)-(4), (6)-(8) & (13) drive growth while *c*(*R*_*N*_) > *c*_*N*_, until the necrotic radius is such that *c*(*R*_*N*_) = *c*_*N*_, at which time the standard Greenspan model holds and growth is driven by Equations (2)-(8).

### 2.3 Summary: statement of full model (dimensionless)

The tumour growth model combined with fractionated radiotherapy (dose *d*_*i*_ Gy delivered at times *t* = *t*_*i*_, *i* = 1, 2,…) can be summarised as follows:

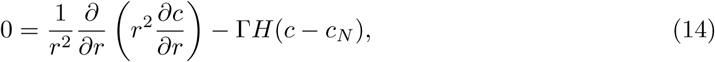

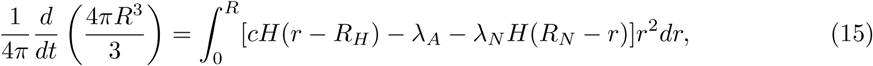

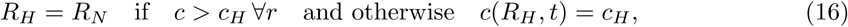

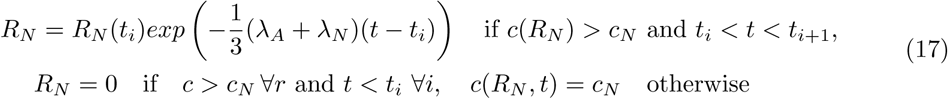

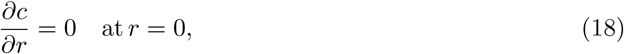

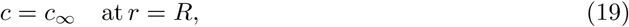

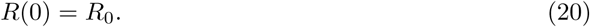

Continuity conditions for *c* and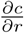 continuous across *r* = *R*_*H*_, *R*_*N*_ are imposed.

Radiotherapy, of dose *d*_*i*_ applied at times *t* = *t*_*i*_, effects an instantaneous volume change. The survival fraction of the normoxic tumour cell population is given by

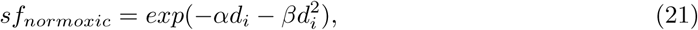

whilst the survival fraction in hypoxic regions is given by

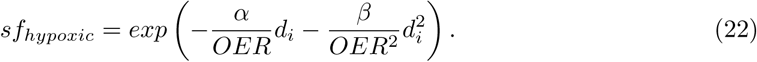

The necrotic core of the tumour spheroid is unaffected by radiotherapy. Immediately after a dose of radiation, the outer tumour radius is given by

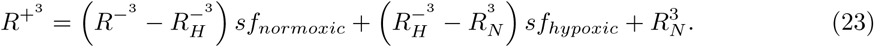

NB: We subtly alter the equation for *R*_*H*_ from the original Greenspan model so that *R*_*H*_ = *R*_*N*_ if the threshold for hypoxia, *c_H_* is not reached. This does not change the dynamics in the case of untreated tumour growth, but ensures that the ordering of the tumour boundaries is conserved in the post-radiotherapy regime (0 **≤** *R_N_* **≤** *R*_*H*_ **≤** *R*).

## 3 Investigation of model behaviour

We now consider the effects of various treatment protocols on a given tumour under this model. In Section 3.1 we solve the model numerically and highlight key features of the simulations for different parameter combinations. We explore these features in more detail in the following sections. We analyse the rapid increase in hypoxia during regrowth of the tumour spheroid post-radiotherapy in Section 3.2. In Section 3.3 we study tumour regrowth following a single dose of radiation, and in Section 3.4 we consider the long-term behaviour of a tumour for different fractionation schedules.

### 3.1 Numerical simulation results

For each tumour growth regime, the oxygen profile *c*(*r, t*) may be solved analytically. Equations (14)-(23) may then be reduced to an ODE for *R* and two algebraic equations for *R*_*H*_ and *R*_*N*_, as described in Appendix A.1. We solve the resulting system of equations numerically using a finite difference scheme implemented in MATLAB.

In Figure 4a we present results showing a typical tumour response to a conventional fractionation schedule (2 Gy, 5 days / week) simulated for 6 weeks using the parameter values in Tables 1 & 2. Cell death induced by the first fraction of radiation delivered at *t* = 1 results in the loss of the hypoxic annulus. While *c*(*R*_*N*_) > *c*_*N*_, the necrotic core decays exponentially, as defined by Equation (13). Figure 5a shows how the oxygen concentration in the necrotic core evolves throughout the first week of the radiotherapy protocol. We see the mismatch between *c*(*R*_*N*_) and *c*_*N*_ following delivery of the first fraction (at *t* = 1). As the tumour grows between fractions, oxygen concentration in the necrotic core decreases, while each fraction of radiation and the corresponding volume loss results in reoxygenation.

**Figure 4:**
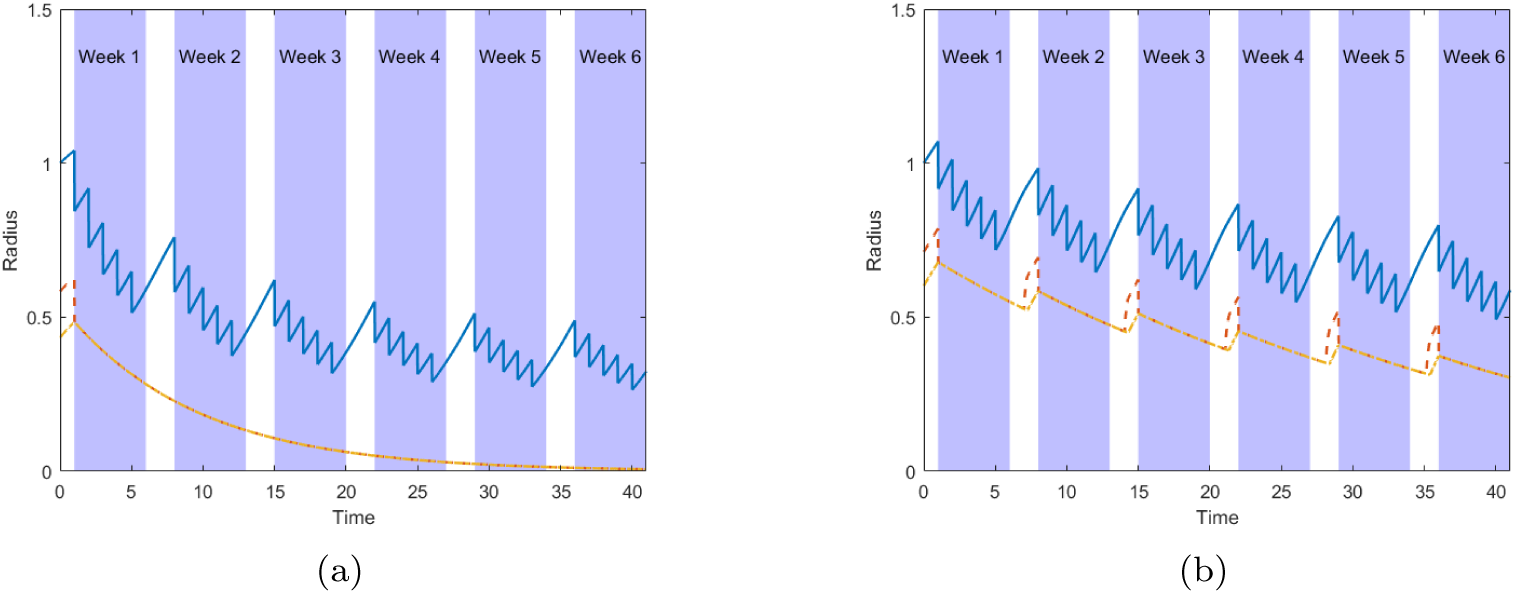
Two solution of Equations (14)-(23) in response to a standard fractionation protocol (2 Gy fractions delivered daily Monday-Friday) simulated for 6 weeks. Overall tumour radius, R blue solid line; hypoxic radius, *R*_*H*_ red dashed line; radius of the necrotic core, *R*_*N*_ yellow dot/dashed line. In (a) we use the parameter values given in Tables 1 & 2 with an initial tumour radius of 3. In (b) we simulate irradiating a tumour spheroid with lower rates of apoptosis (*λ_A_* = 0.1213) and oxygen consumption (Γ = 0.3032) starting from an overall radius of 7.5. The evolution of the tumour contours *R, R_H_* and *R*_*N*_ is shown by the blue, red and yellow lines, respectively.

**Figure 5:**
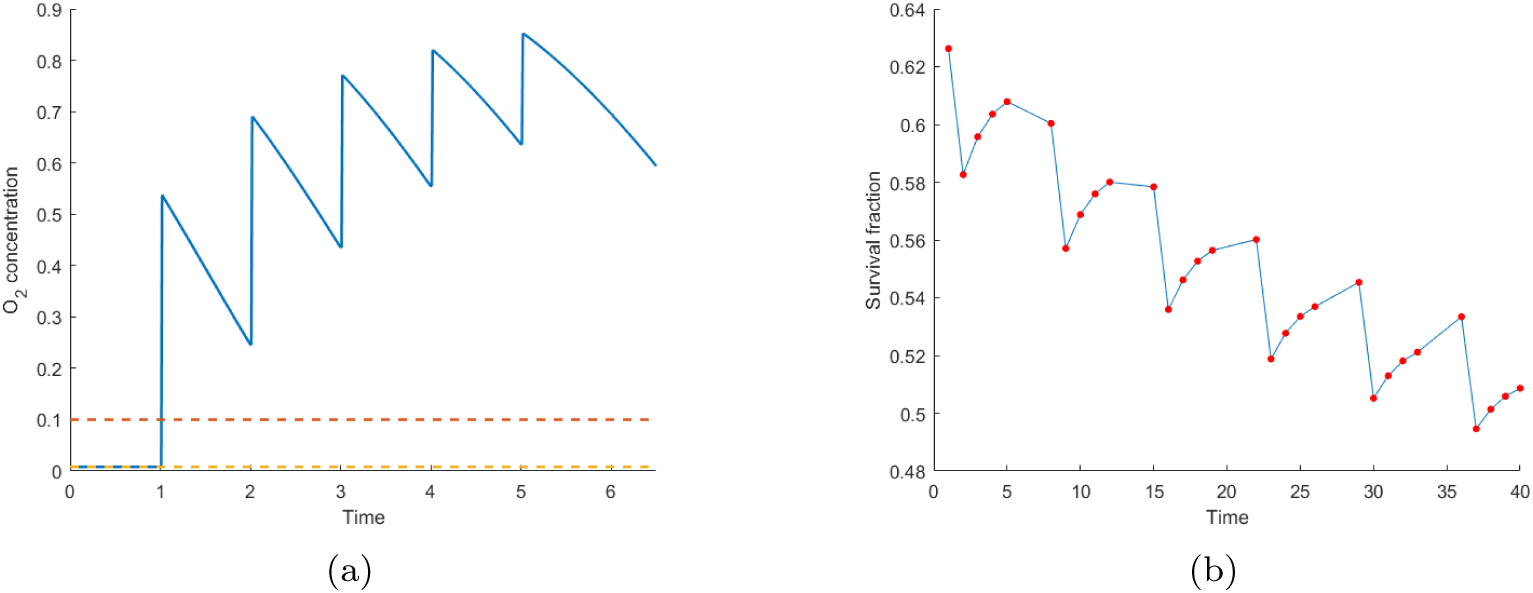
Two plots highlighting key behaviours in the simulations shown in Figure 4. (a) Blue line shows oxygen concentration within the necrotic core during the first week of treatment for the simulation in Figure 4a. The oxygen thresholds for hypoxia, *c_H_*, and necrosis, *c*_*N*_, are shown by dashed red and yellow lines, respectively. (b) Change in average tumour survival fraction following successive doses of radiotherapy corresponding to the simulation presented in Figure 4b. Red points correspond to the survival fraction for each fraction of radiotherapy delivered, blue line added for ease of visualisation. (NB. y-axis not [0,1])

For this parameter combination, radiotherapy results in a tumour volume at the end of treatment that is sufficiently small such that the spheroid is almost entirely composed of proliferative cells. In particular, the necrotic core has decayed so that *R*_*N*_ ≪ *R*.

Figure 4b shows how the system dynamics change when a tumour with a much lower apoptotic rate (*λ_A_*) is exposed to the standard fractionation protocol. The slower rate of apoptosis results in a larger steady state tumour in the absence of radiotherapy. Similarly, a larger tumour volume supported throughout treatment and a more gradual volume reduction. In this case the hypoxic annulus reappears during treatment (as observed during the weekend breaks in the protocol) since the necrotic core decays such that *c*(*R*_*N*_) = *c_H_* before the end of the fractionation protocol. A rapid transient increase in *R*_*H*_ is observed when the hypoxic annulus reappears. We characterize this behaviour in Section 3.2.

For the simulation in Figure 4b, a tumour spheroid comprising a necrotic core and transient hypoxia persists at the end of treatment. The tumour volume appears to be evolving towards a periodic orbit and, as such, the model predicts that continuing the same fractionation schedule indefinitely will not yield significant further volume reduction.

Since the tumour composition changes dynamically throughout treatment, the efficacy of each radiation fraction also varies. The survival fraction changes by about 10% from the start to the end of treatment due to the shrinkage of the necrotic core (Figure 4b). In this case a characteristic shape within the curve in Figure 5b repeated weekly due to the weekly oscillations in tumour composition, with the spike associated with the rapid reappearance of hypoxia. Such dynamic behaviour clearly depends on tumour-specific parameters and highlights the additional details that can be observed when spatial effects are included in a mathematical model (c.f. difference in tumour composition between Figures 4a & 4b).

### 3.2 Asymptotic analysis of model behaviour post-radiotherapy

When a fully-developed tumour spheroid including a hypoxic annulus and necrotic core is exposed to fractionated radiation, the composition of the spheroid immediately after irradiation depends on the dose *d*, as described in Section 2.2.1. If the dose is sufficiently large, the post-radiotherapy composition comprises a proliferating rim and a necrotic core, but with the absence of a region of hypoxia. In this scenario, the oxygen concentration in the necrotic core is greater than the threshold for hypoxia (i.e. *c*(*R*_*N*_) > *c_H_*). Upon regrowth, the outer tumour radius, *R*, will increase, while *R*_*N*_ will decrease as the necrotic core undergoes decay. Since this parameter regime yields tumour spheroids with proliferating, hypoxic and necrotic compartments prior to radiotherapy, left untreated the tumour will eventually evolve so that the hypoxic annulus will start to re-develop (when *c*(*R*_*N*_) = *c_H_*). This sequence of events is summarised in Figure 6. Simulation results reveal the re-emergence of hypoxia following radiotherapy via a fast, transient increase in *R*_*H*_ (see Figure 4b). Since hypoxic cells are less radio-sensitive, such a rapid increase in the hypoxic volume could have implications for the response to further doses of radiation. We now characterise this behaviour.

**Figure 6:**
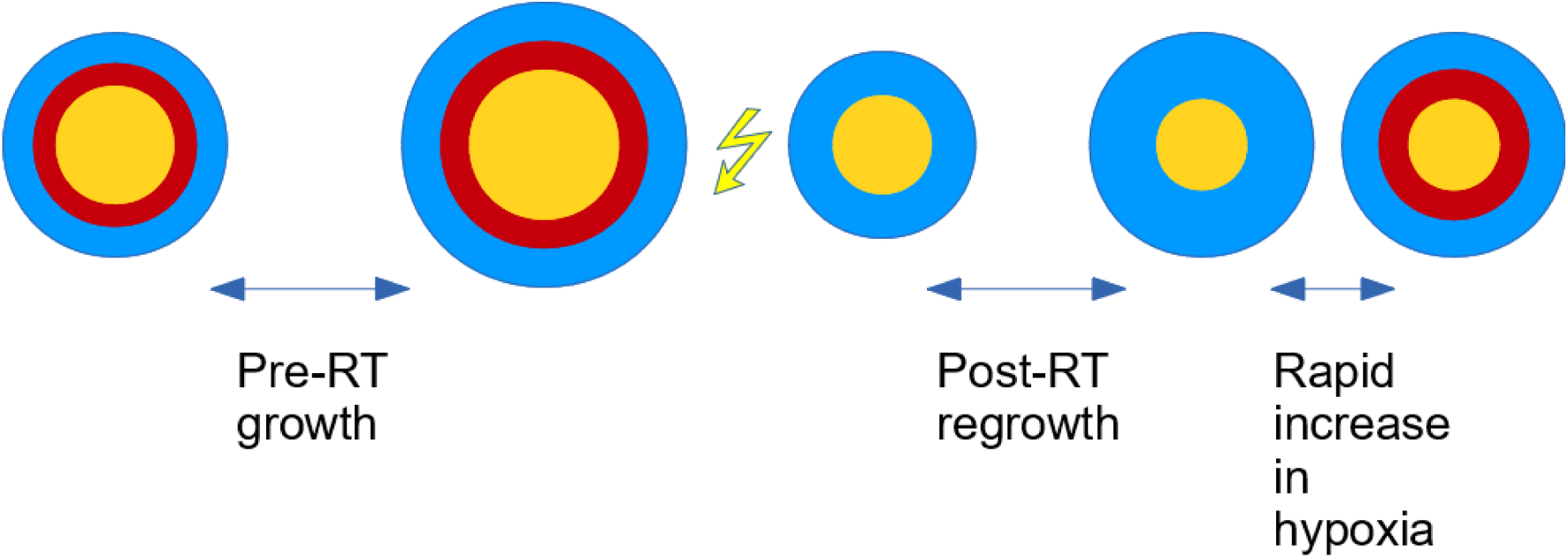
Schematic contextualising the scenario in which the rapid transient increase in hypoxia is observed, corresponding to the different growth phases observed in Figure 7.

**Figure 7:**
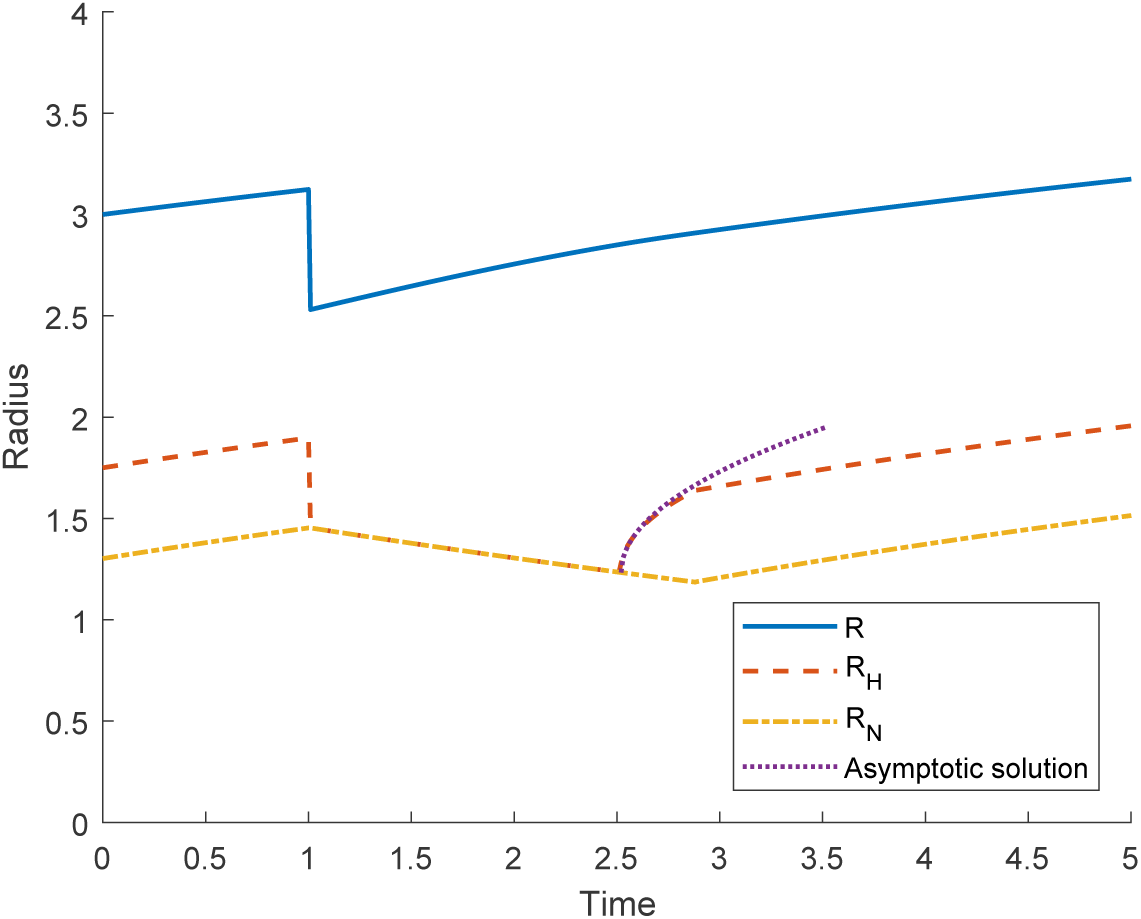
Asymptotic solution given by Equation (38) (dotted line) plotted alongside the numerical solution to Equations (14)-(23) showing the regrowth of the tumour spheroid after radiotherapy and the fast initial increase in *R*_*H*_. We note that during regrowth the mismatch in the oxygen tension on the necrotic core boundary is eventually resolved and the tumour resumes standard Greenspan growth dynamics.

The fast initial increase in *R*_*H*_ upon regrowth of the tumour spheroid in the Greenspan model after a radiation fraction (see Figure 7) occurs when the necrotic core is undergoing exponential decay and a region of hypoxia is about to develop outside the necrotic core. We analyse this situation by supposing that (without loss of generality at *t* = 0) *c*(*R*_*N*_) = *c_H_*. Then solution of Equations (14), (18) and (19) yields

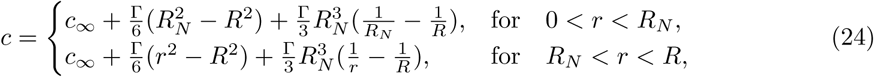

while Equation (15) reduces to give

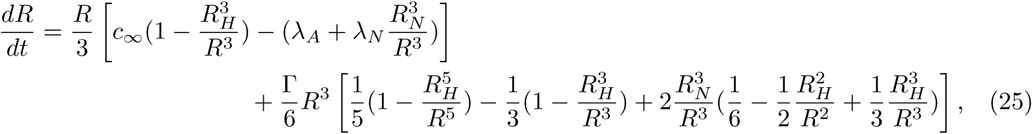

and the internal free boundaries *R*_*H*_ (*t*) and *R*_*N*_ (*t*) satisfy

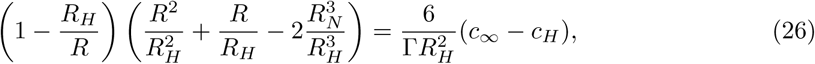

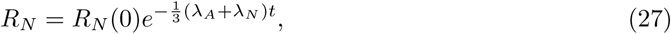

with *R*_*H*_ (0) = *R*_*N*_ (0), and *R*(0) defined by Equation (26).

With *R*_*N*_ (*t*) defined by Equation (27), we seek approximate solutions for *R* and *R*_*H*_. Differentiating Equation (26) with respect to t we obtain an ODE for *R*_*H*_ (*t*), which is singular in the limit as *R*_*H*_ → *R*_*N*_:

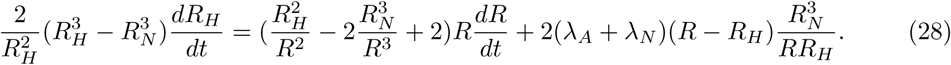

We investigate the behaviour in this limit by making the change of variables 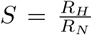with *S* = 1 at *t* = 0. Then Equations (25) and (28) become

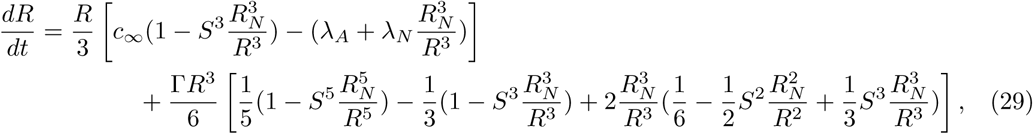

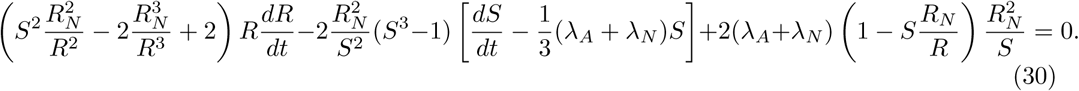

We construct approximate solutions to Equations (29) and (30) in the limit when *S* is close to 1. Introducing the small parameter ϵ (0 < ϵ ≪ 1), we consider the short timescale *t* = *ϵ*^2^*τ* and propose expansions for *S* and *R* in this boundary layer of the form

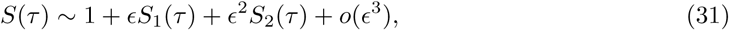

and

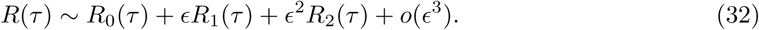

The choice of timescale can be justified by a dominant balance argument, allowing us to regularise the ODE for *S* at leading order. Note that *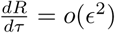*, so we deduce that

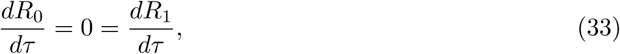

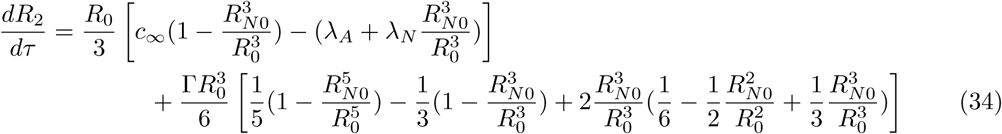

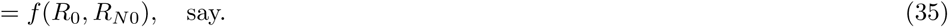

Hence, R ∼_ R_0_ + ∊^2^R_2_(τ) + o(∊^3^),where *R*_0_ = *const* = *R*(0). We also find that *R*_*N*_ has no *o*(*ϵ*) term, since expanding Equation (27) gives 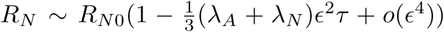 where *R*_*N*_0 = *R*_*N*_ (0).

Turning our attention back to Equation (30), we find at leading order

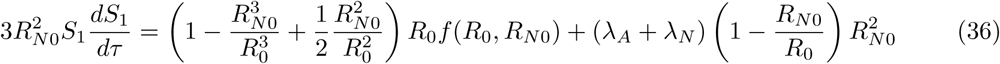

where *f* (*R*_0_, *R*_*N*0_) is defined via Equation (35). Integrating Equation (36) gives

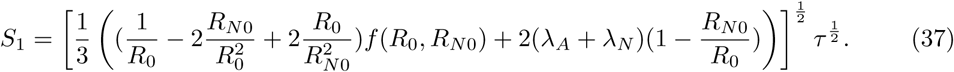

So for *t* **≪** 1,

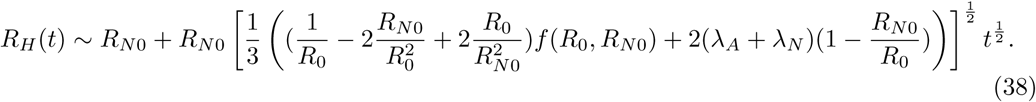

This approximate solution for *R*_*H*_ (*t*) and the numerical solution obtained by solving the full problem are in good agreement (Figure 7).

As a result of this analysis we conclude that whenever a hypoxic region re-emerges within a tumour post-radiotherapy, it does so rapidly over a short timescale on which both the outer tumour radius and the radius of the necrotic core do not change dramatically. This phenomenon arises naturally from the model. We note that similar analysis was performed on the original, untreated growth equations by Byrne *et al.* (Byrne and Chaplain, 1998).

### 3.3 Single hit regrowth

We now consider the effect of irradiating a small tumour spheroid composed entirely of proliferating cells and its subsequent regrowth. This situation is relev t forantumours of radius *R* such that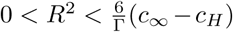and *R*_*H*_ = *R*_*N*_ = 0, since when *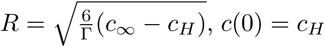*and thereforelarger tumours will contain a hypoxic region and/or necrotic core. Solving Equation (14) subject to boundary conditions (18)-(19) we deduce that the oxygen profile for this tumour composition is given by

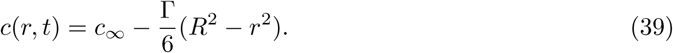

Substituting from Equation (39) into Equation (15), with *R*_*H*_ = *R*_*N*_ = 0, we arrive at the following ODE for *R*(*t*):

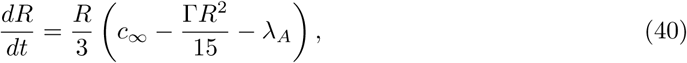

with solution

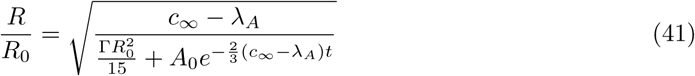

where *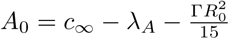*

We can use Equation (41) to calculate the time taken for a tumour spheroid of radius *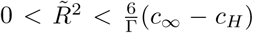*following a single fraction of radiation of dose *d* to regrow to its initial (i.e. pre-radiotherapy) size. The tumour radius immediately after irradiation is given by 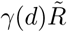 where *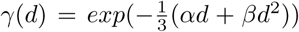*is the survival fraction given by the linear quadratic formula. It is straightforward to show that the time, Δ *t*, until the tumour regrows to its original size is given by

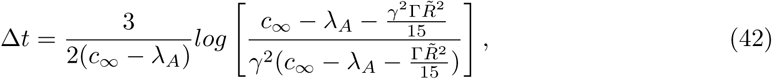

where *γ* = *γ*(*d*). We assume that the parameters are such that the tumour spheroid was growing pre-irradiation and therefore 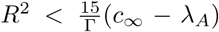 (from (40)). In this parameter regime, the logarithm in Equation (42) is defined and Δ *t* > 0. By extension, we require that *c _∞_*> *λ_A_* since otherwise the tumour spheroid shrinks for all values of 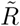 and a viable tumour of any size cannot be supported. We note for future reference that

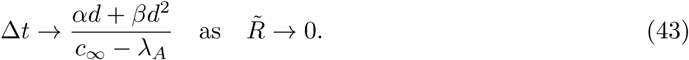

Differentiating Equation (42) with respect to 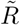, we obtain

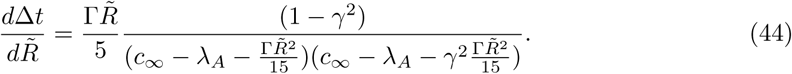

Therefore,at least for small 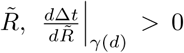, and so the regrowth time, Δ *t*, is an increasing function of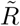 for fixed dose *d*. This holds for all 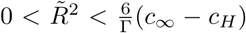if the inequality 3c_∞_ – 5λ_*A*_ + 2*_cH_* > 0 is satisfied. Then Equation (42) represents an increasing family of curves where the minimum bounding curve is given by Equation (43). That is, for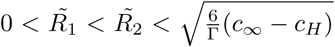and fixed dose *d*, Δ*t*(*d*;*R*_1_) < Δ*t*(*d*;*R*_2_)

Returning to Equation (42), we see that for an initial radius 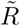a strategy that combines a dose *d* with an inter-fraction time less than Δ *t* will result initially in a net shrinkage of the tumour throughout treatment (i.e. the tumour volume is smaller at the delivery of each radiation fraction). Conversely, if we wait longer than Δ *t* to re-irradiate then the tumour will have grown larger than its original size. Thus, Equation (42) defines a curve in the (*d,* Δ *t*)-plane for treatment protocols which give rise to periodic behaviour for a given initial tumour radius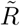since the cell kill induced by radiotherapy is exactly balanced by the regrowth between fractions. This curve is indicated in Figure 8. We note that the curve describing protocols that yield periodic behaviour becomes concave for large *R*, and in particular, as *R* tends to its steady state value, *R\** say, then for a given dose, *d* > 0, Δ *t*(*d*) *→ ∞*.

**Figure 8:**
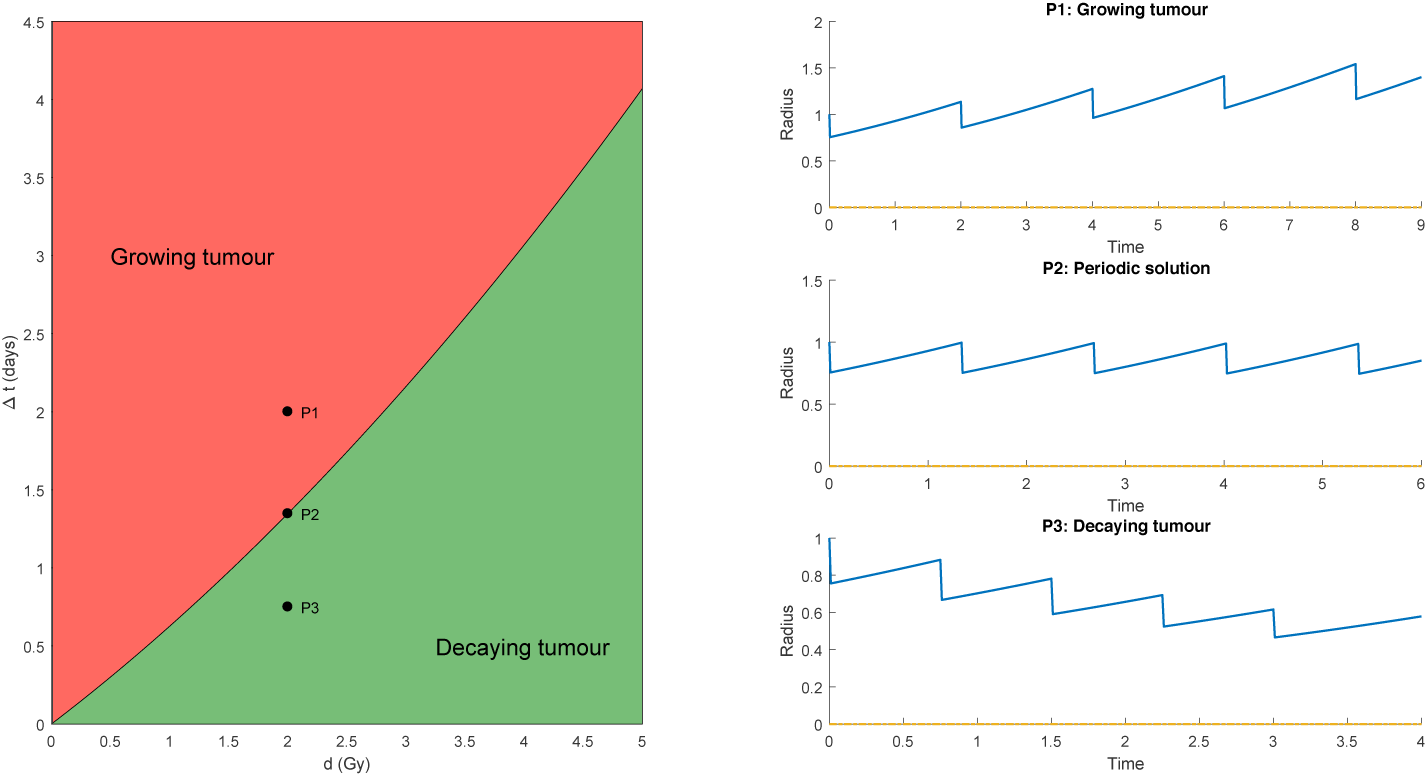
Radiation dose and fractionation-dependent behaviour for a tumour spheroid composed of proliferating cells. Line for periodic behaviour given by Equation (42). Plots on the right show response to 5 fractions of radiotherapy given by the protocols corresponding to points P1, P2 & P3.

### 3.4 Periodic surface in ‘treatment space’

In the previous section we investigated the response of tumours of fixed initial radius 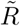to different fractionation protocols as specified by a dose, *d*, and inter-fraction time, Δ *t*. The (*d,* Δ *t*)-plane can be partitioned into regions in which the tumour volume increases or decreases during treatment. However, we note that the location of these regions depends on the tumour radius at the time of irradiation, 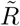As such, when considering the behaviour of a tumour throughout an entire course of radiotherapy, it is not a line in the (*d,* Δ *t*)-plane that we are interested in, but a surface in ‘treatment space’, Ͳ ⊂ ℝ^3^, with components (*R, d,* Δ *t*).

For a protocol delivering *n* fractions of radiation, we can identify the response of a given tumour with a discrete trajectory, (*R*(*t*_*i*_)*, *d*_*i*_,* Δ *t*_*i*_) ∈ Ͳ, through treatment space. Note, we consider the trajectory as discrete points pre-irradiation so that *R*(*t*_*i*_) = *R*(*t*_*i*_)- in our previous notation. Here *t*_*i*_, *d*_*i*_ and Δ *t*_*i*_ are the time, dose and inter-fraction time, respectively, of the *i*th fraction, for*i* = 1,…, *n*. The union of the curves in the (*d,* Δ *t*)-plane determining periodicity define a surface, S, in T (see Figure 9). Now, as a tumour progresses through the course of radiotherapy, points on its trajectory that lie above this surface indicate net growth of the tumour by the time of delivery of the next radiation dose, whilst points below the surface result in a more desirable decrease in tumour volume. Therefore, for a given tumour spheroid, S defines a surface that partitions T into treatment protocols that cannot halt tumour progression and those that lead to tumor decay.

**Figure 9:**
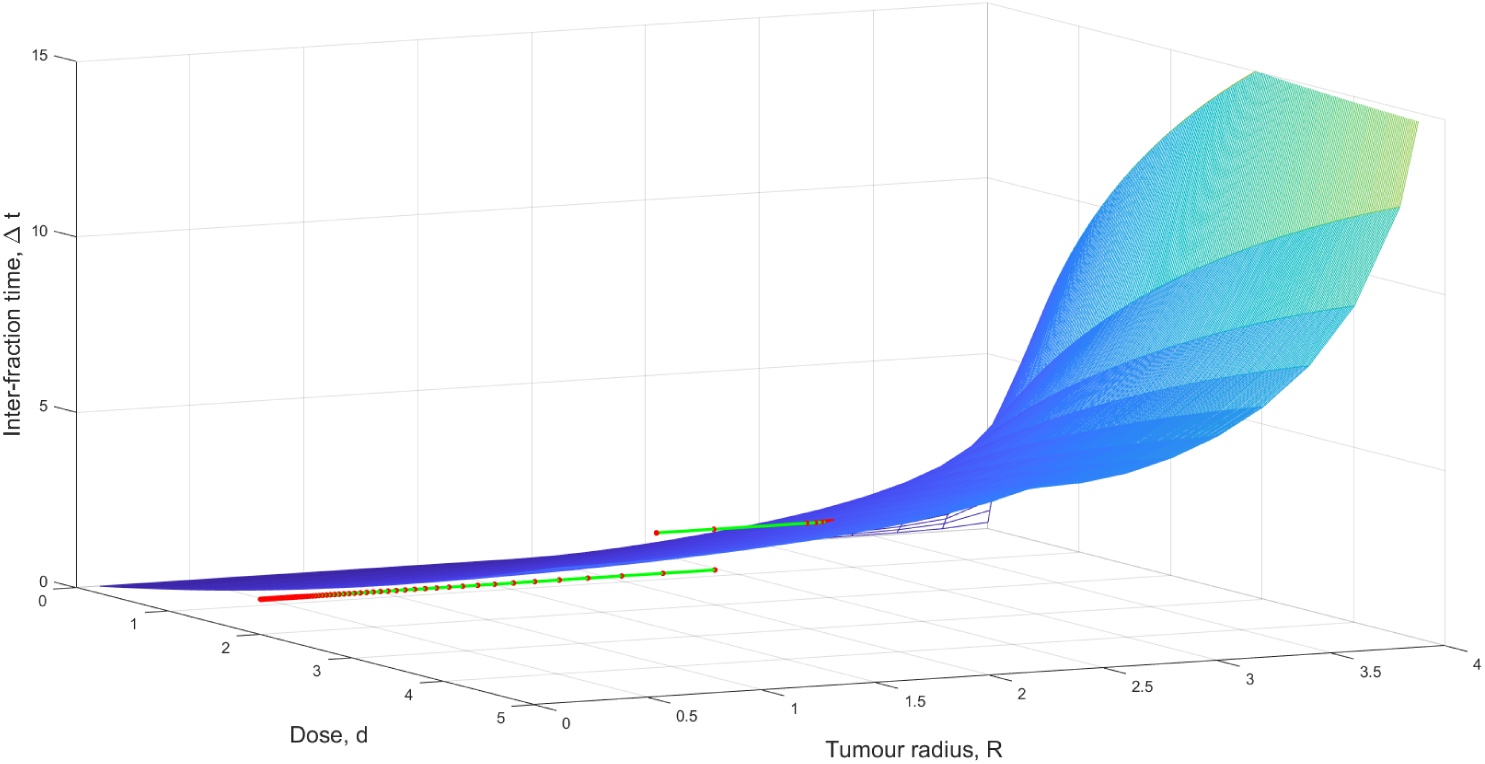
Simulated trajectories (green lines) in T for two different constant dosing schedules (fractions plotted as red points) applied to the same tumour spheroid (i.e. same model parameters). A dose of 3 Gy every 3 days (top trajectory) results in a tumour that grows until it reaches a periodic state, while a dose of 2 Gy delivered daily (bottom trajectory) is sufficient to drive the tumour volume arbitrarily close to zero. Surface describes treatment protocols which give rise to periodic behaviour.

With this understanding more general statements about the outcome of different fractionation protocols can be made. We first consider dosing schedules in which the same dose *d* is administered at constant intervals Δ *t*. In this case, if a point in T lies above S, then the tumour spheroid will grow until it converges to a point on S, at which time it becomes periodic (Figure 9).

Below S, then the outcome of radiotherapy depends on the boundary curve of S as *R* → 0 (see Equation (43)). If the fractionation schedule lies above this curve in the (*d,* Δ *t*)-plane then the system will eventually converge to the corresponding point on S. However, *d* and Δ *t* below this can be chosen such that the tumour decays to arbitrarily small volumes (Figure 9). We note that Equation (43) still holds for this analysis even though its derivation requires an initial tumour composed entirely of viable cells. This is the case since, in the model, necrotic material is only created via nutrient starvation and not as a result of irradiation. As such, for ongoing treatment protocols that result in a decreasing tumour volume, as *t → ∞, R* **≫** *R_N_*.

We can extend this concept to more general treatment schedules. If there exists *m* **∈;** ℕ such that for all *i* > *m* the point (*di,* Δ *ti*) of the *i*th fraction lies below the curve given by Equation (43), then treatment is sufficient to drive the tumour volume arbitrarily close to 0. However, using Figure 4b as an illustrative example, we observe a simulation in which the weekend break of the standard fractionation protocol corresponding to Δ *t* = 3 days gives rise to the periodic orbit.

When predicting how a tumour may respond to radiotherapy, two quantities of interest are the *potential volume doubling time*, *Tpot*, and the *survival fraction after 2 Gy of radiation*, *SF*2, of the tumour. We translate these definitions into the model and identify tumour characteristics for which this model would eventually predict regrowth over the weekend.

The survival fraction at 2 Gy, *SF*_2_, for normoxic tumour cells is given by the linear-quadratic formulation. In order to find an expression for *T*_*pot*_, we consider the tumour dynamics for small tumour volumes that are in an ‘exponential growth‚ phase. For small R,

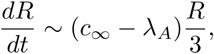

which yields a volume doubling time of

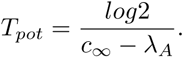

Substituting these expressions into Equation (43) for a standard 2 Gy dose we obtain

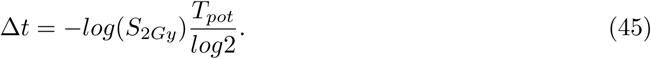

We note that this expression holds for tumour growth models in which the growth of small tumours is exponential and radiation-induced cell death is modelled as an instantaneous volume loss.

If we now consider Equation (45) in the context of the weekend break (Δ *t* = 3) within the standard fractionation protocol, then we see that -*T*_*pot*_*log*(*S*_2*Gy*_) < 3*log*2 defines a class of tumour characteristics for which regrowth of the tumour over the weekends impedes tumor decay (Figure 10).

**Figure 10:**
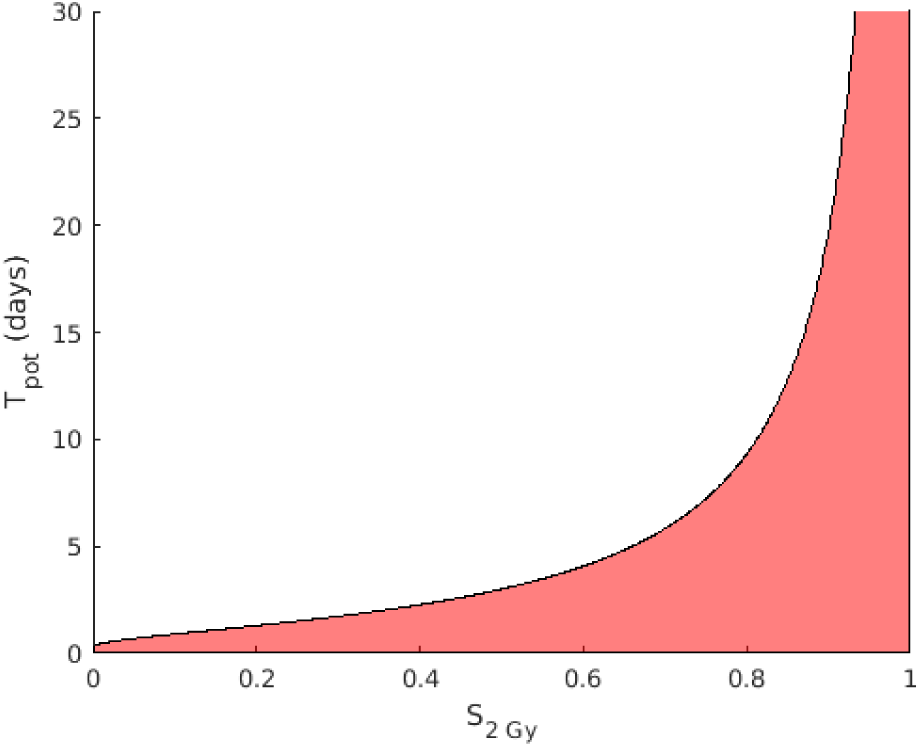
Region for which tumours will eventually exhibit net growth over the weekend (shown in red).

## 4 Discussion

*In vivo* tumours are highly heterogeneous cellular entities characterised by high inter-patient variability. Allied to this, the local efficacy of radiotherapy delivered to the tumour site is affected by a number of variables associated with the tumour’s heterogeneous composition and microenvironment. In particular, the local oxygen concentration can significantly influence radiation-induced cell death, with well-oxygenated regions being shown to exhibit up to threefold greater radioensitivity than hypoxic tumour populations. With this in mind, in this paper we have presented a simple, spatially-resolved model in order to investigate the effects of tumour composition on radiotherapy response. The discussed model builds on the tumour growth model first proposed by Greenspan (Greenspan, 1972). We extend this to incorporate the effects of radiotherapy taking into account spatially-varying radiosensitivities.

Numerical simulations and mathematical analysis of the model reveal how the tumour’s growth dynamics and spatial composition change throughout treatment. Heterogeneity within the tumour not only affects the initial response to radiotherapy, but also how this response changes throughout the duration of the treatment protocol (see Figure 5b). For parameter regimes in which hypoxia reemerges during treatment, a rapid transient increase in the width of the hypoxic annulus that is observed. This behaviour arises naturally from the model and the underlying process driving this phenomenon remains to be elucidated. The model more generally also classifies protocols that may result in tumour progression, a non-zero periodic tumour volume, or tumor decay. We identify a surface in ‘treatment space‚ dependent on tumour-specific growth and radio-sensitivity parameters and determine that successful protocols correspond to those that remain below this surface throughout treatment. The wide variety of dynamics observed suggests that spatial heterogeneity may be important for simulating tumour response to radiotherapy and, in particular, for making clinical predictions.

The model presented in this paper makes numerous assumptions and simplifications about the underlying biology. For radiation-induced cell death we take the common approach of modelling this process as an instantaneous effect. However, the linear-quadratic model was established to determine long-term clonogenic survival after radiotherapy. Biologically, cell death after radiation may occur via a number of different mechanisms, with many irradiated cells dying only after attempting mitosis one or more times (Joiner and van der Kogel (2009), Chapter 3). Consequently, radiation-induced cell death may not elicit the instantaneous volume reduction modelled here. In future work we will model this process in more detail in order to describe the short term response to radiotherapy and the corresponding spatial changes in tumour composition more accurately. Additionally, a more detailed model will allow us to relax the restrictions of the spherically symmetric geometry associated with the current model, making it more applicable to modelling vascular tumour growth and *in vivo* responses to radiotherapy.

However, for a model to be useful in making patient-specific predictions in the clinic, parameters must be identifiable with respect to the limited clinical data typically available. As such, current research in this area often focusses on phenomenological ODE models with few parameters to be estimated. Typically these models contain no information about the spatial heterogeneity in tumour composition and radiotherapy response. With this in mind, we also propose a compar-ison of the developed spatially-resolved model with these phenomenological approaches as future work. We aim to identify situations in which the spatially-resolved and spatially-averaged models agree well, and those in which there is a significant difference. The tumour composition changes observed in model simulations suggests that averaged parameter values in simple, phenomenological models may not sufficiently capture the tumour dynamics during treatment for some tumour compositions and parameter combinations.

While more sophisticated models may be difficult to parametrise in practice, they have the potential to increase biological insight and inform further modelling studies. Spatially-resolved models, such as the one presented in this paper and the future work proposed to generalise some of the simplifying assumptions made here, may aid in the development of alternative clinically-focussed models which capture more of the key features than existing phenomenological models. More complex models incorporating more biological detail may also be used to generate data for *in silico* testing of ODE models and model selection paradigms; comparing the quality of fit and future predictions of a range of simple models against a known‘ground-truth**‚**.

## 5 Acknowledgements

This work was supported by the Engineering and Physical Sciences Research Council [grant number EP/G037280/1]. TL would also like to thank the Moffitt Cancer Center, where some of this work was undertaken, for their hospitality.

## A Appendix

### A.1 Explanation of Greenspan’s original growth model

Here we provide further explanation of Greenspan’s original model for the nutrient-limited growth of tumour spheroids (Greenspan, 1972). We restate the equations for the evolution of the nutrient concentration profile, *c*, and the tumour radii, *R*, *R*_*H*_ and *R*_*N*_.

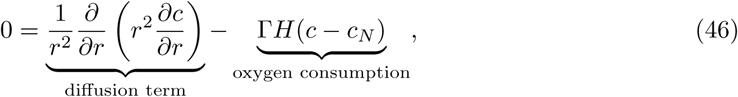

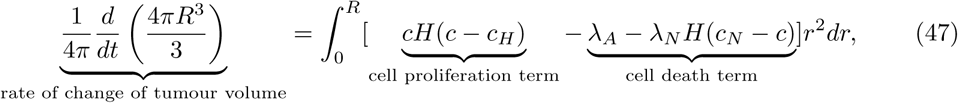

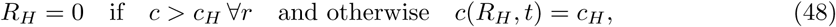

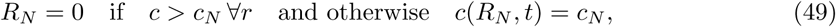

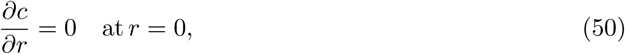

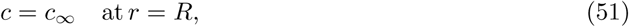

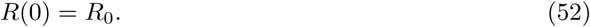

In (46), we assume that the oxygen concentration within the tumour is regulated by its diffusion across the tumour (with diffusion constant *D*) and consumption by the tumour cells. We make the further assumption that both normoxic and hypoxic cells consume oxygen at the same constantrate, Γ, and that cells in the necrotic core do not consume oxygen. Since oxygen diffuses on a much shorter timescale than tumour growth we assume that the oxygen concentration is in a quasi-steady state. Hence, *c*(*r, t*) satisfies (46), where *H*(.) is the Heaviside function, with associated boundary conditions (50) and (51) imposing symmetry at *r* = 0 and a Dirichlet boundary condition at *r* = *R*(*t*).

We assume that rates of cell proliferation and death within the tumour are determined by the local oxygen concentration. Proliferation occurs at a rate proportional to *c* where there is sufficient oxygen supply (*c* > *c_H_*). Both apoptosis and necrosis contribute to cell death within the tumour, with apoptosis occurring at a constant rate throughout the tumour and necrosis localised to the necrotic core (where *c* **<** *c*_*N*_). Degradation of the necrotic core is assumed to result in material that is freely permeable throughout the tumour spheroid. We assume that adhesion and surface tension forces acting on the tumour cells maintain the shape of the tumour spheroid, and that these same forces push cells inwards to compensate for the outward flux of necrotic material from the necrotic core. The evolution of *R*(*t*) is then given by the mass balance in Equation (47), with *λA*, *λN* > 0, accompanied by the initial condition (52). The radii at which the tumour becomes hypoxic and necrotic, *R*_*H*_ and *R*_*N*_, respectively, are determined as contours of the oxygen concentration profile (Equations (48) and (49)). For situations in which the tumour spheroid is small enough that one or both of these contours do not exist, we define the corresponding radius to be 0.

An analytic solution can be found for *c*(*r, t*) that depends on *R, R_H_* and *R*_*N*_, and so this system can be reduced to an ODE for *R* and algebraic equations for *R*_*H*_ and *R*_*N*_. The resulting equations in the case of a fully developed, 3-layer tumour spheroid are given in Equations (53)-(57). The system of equations for the earlier growth phases are similar and can be obtained in the same manner.

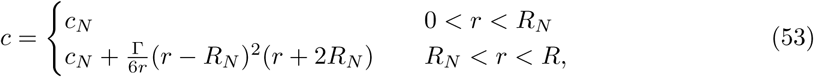

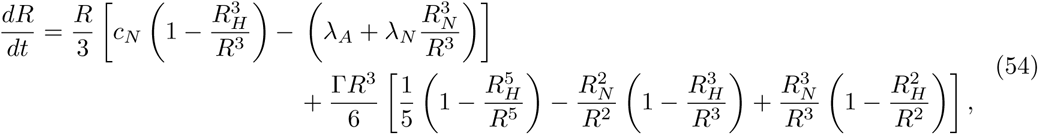

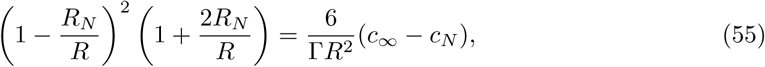

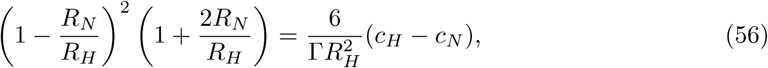

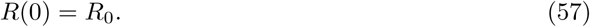

Figure 2 (main text) demonstrates the growth and various stages (labelled C1, C2 & C3) of tumour spheroid composition under the Greenspan model using the parameter values in Table 3. In using partial pressures of oxygen rather than concentration, we follow Grimes *et al.* (Grimes et al, 2014) and use Henry’s law to convert between the two, so that *p* = Ω*c*, with Ω = 3.0318x107 mmHgkgm^-3^. The oxygen concentration profile at each time point is a monotonic function increasing outwards from the tumour centre. Initially, for very small, avascular tumour spheroids,the entire tumour is made up of viable, proliferating cells (case A). We notice that as the tumour grows and so the oxygen concentration at the centre of the tumour decreases, a hypoxic region within the tumour begins to develop (case B) followed by a central necrotic core when the oxygen concentration falls so low as to be unable to support viable cells (case C). These different regions, and their varying responses to radiation, require careful consideration when extending Greenspan’s model to account for radiotherapy. Further discussion of the different phases of growth is given in (Byrne, 2012).

**Table 3:** Parameter values for Greenspan’s model of tumour spheroid growth. (Note: dimensionalapoptosis and necrosis rates, *s-*^1^, are given by *s λ_A_* and *s λ_N_*, respectively. *p _∞_*, *p*_*H*_ and *p*_*N*_ are partial pressures which, for consistency, are then converted into concentrations *c ∞*, *c_H_* and *c*_*N*_, respectively. For further details see Appendix A.1.)

| Parameter | Symbol | Value | Source |
| --- | --- | --- | --- |
| Oxygen consumption rate, $m^3 kg^{-1} s^{-1}$ | $\Gamma$ | $7.29 \times 10^{-7}$ | Grimes et al (2014) |
| Partial pressure at tumour boundary, $mmHg$ | $p_\infty$ | 100 | Grimes et al (2014) |
| Hypoxic partial pressure, $mmHg$ | $p_H$ | 10 | Grimes et al (2014) |
| Anoxic partial pressure, $mmHg$ | $p_N$ | 0.8 | Grimes et al (2014) |
| Oxygen diffusion constant, $m^2 s^{-1}$ | $D$ | $2 \times 10^{-9}$ | Grimes et al (2014) |
| Apoptosis constant, $m^3 kg^{-1}$ | $\lambda_A$ | $1.0555 \times 10^{-6}$ | Frieboes et al (2007) |
| Necrosis constant, $m^3 kg^{-1}$ | $\lambda_N$ | $2 \times 10^{-8}$ | Schaller and Meyer-Hermann (2006) |
| Cell birth/death constant, $kg m^{-3} s^{-1}$ | $s$ | 3.509 | Frieboes et al (2007) |

### A.2

Parameter sweep of Greenspan growth dynamics

We initially investigate the sensitivity of the standard Greenspan growth dynamics to some of its key parameters. In particular, as we sweep over a region of parameter space, we observe the resulting long-time, steady-state properties of the tumour spheroids Specifically,weinvestigatehow the availability of oxygen, *c _∞_*,theoxygen consumption rate of the cells, Γ, and the rates of apoptosis and necrosis, *λ_A_* and *λ_N_*, respectively, affect the growth of the tumour under this model.We choose appropriate intervals for each parameter, in each case ncompassing the corresponding value shown in Table 3, and systematically explore the resulting region of parameter space, solving the model numerically. We plot heatmaps in parameter space in order to observe the behaviourof a variety of quantities of interest of the tumours at steady state (Figures 11 & 12). We have omitted here the results ofvarying both *c _∞_* and *λ_N_* since, in the case of *c _∞_*, increasing the parameter value simply has the effect of anincrease in overall tumour size and the width of thecorresponding proliferating rim as might be expected, while, for the range of values swept over, *λ_N_* had little effect on the observed tumour characteristics.Broadly speaking, we see in Figure 11 that relatively increasing the amount of oxygen available to the tumour cells (by decreasing the consumption rate Γ), or decreasing the death rate, *λ_A_*, results in larger steady state tumours, as we might expect. Point A marks the location of the parameter values from Table 3. Extreme points in the region of concern in parameter space are labelled B, C and D and the corresponding tumour evolution profiles shown. We see that the greater relative availability of oxygen and the low death rate at point B results in a large, fullydeveloped tumour spheroid, whereas the high rate of apoptosis at D is such that a viable, tumour cell population cannot be sustained.

**Figure 11:**
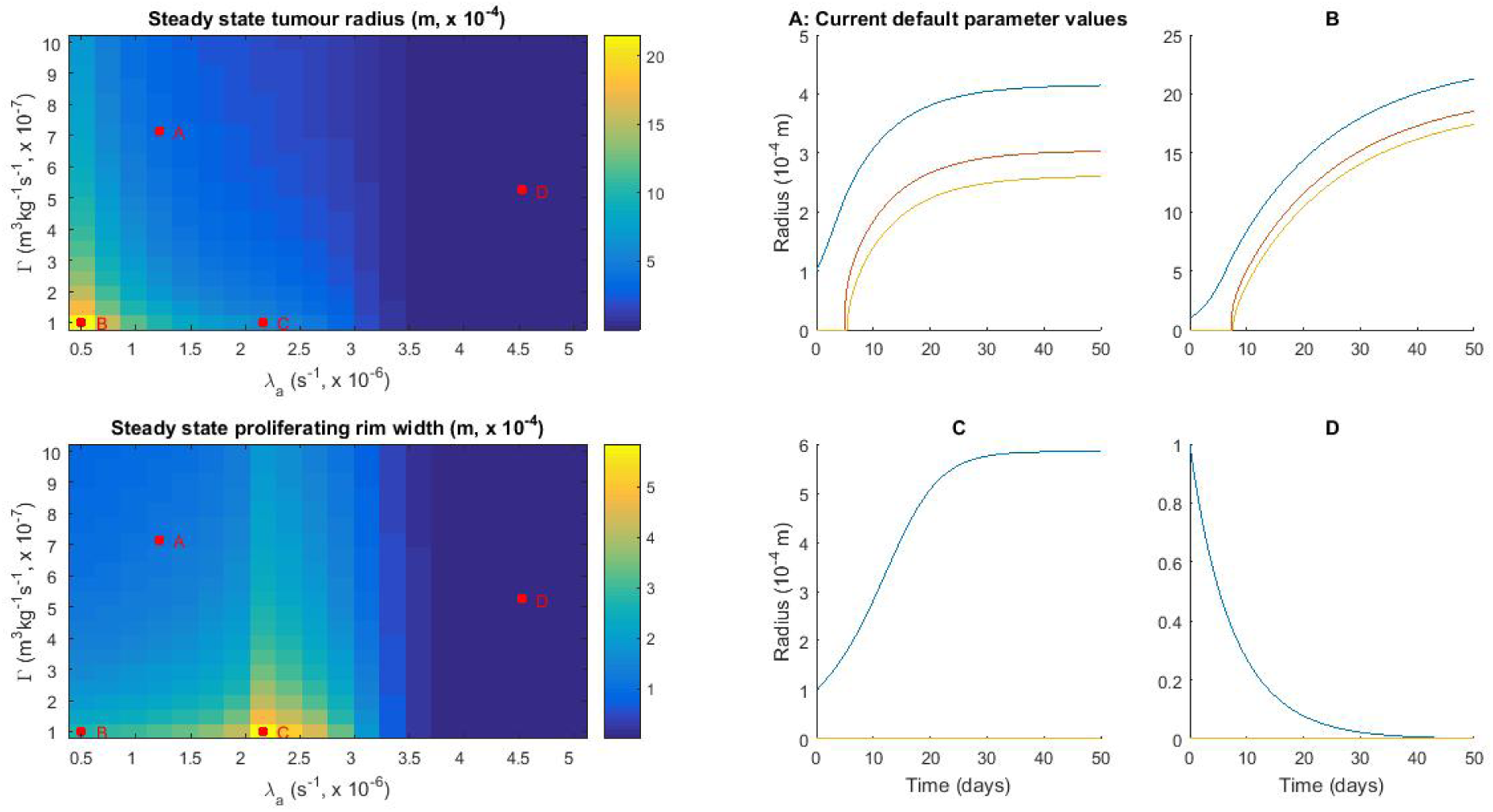
Results of a parameter sweep for the steady state tumour radius and width of proliferating rim with corresponding tumour growth profiles at points of interest in parameter space, A, B, C and D.

**Figure 12:**
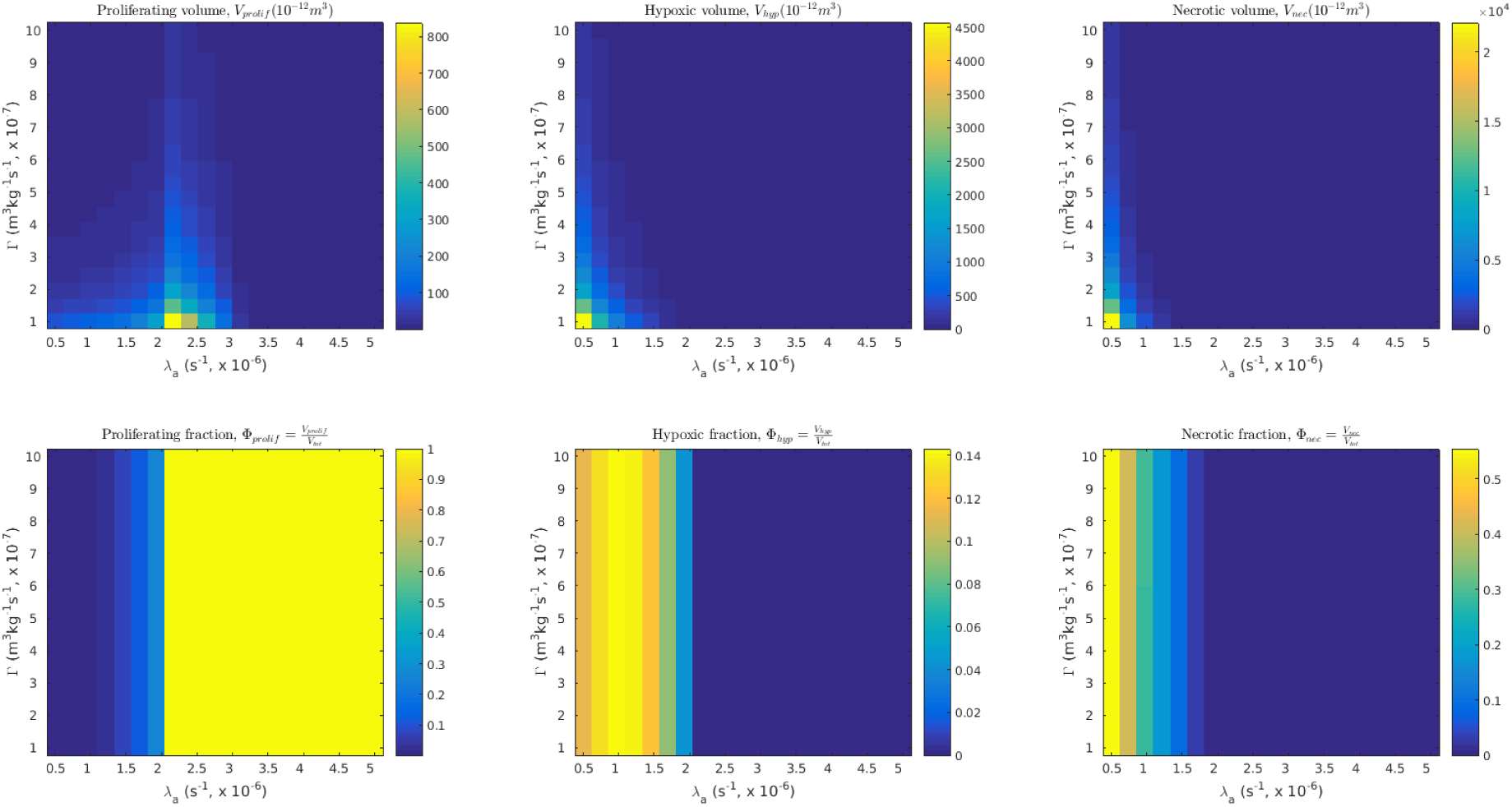
Heatmaps for the steady state volumes of proliferating, hypoxic and necrotic cells, and the corresponding volume fractions.

For a given Γ, we can determine the onset of necrosis as the radius 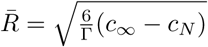At the point C, and any other point on a vertical line through C, the rate of apoptosis is such that the tumour reaches a steady state of radius 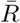and is given by *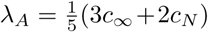*Beyond C, at lower death rates (eg B), we enter a new phase of growth that includes hypoxia and necrosis. The rapid,transient increase in *R*_*H*_ /*R*_*N*_ at onset of hypoxia/necrosis (c.f. asymptotics in Section 3.2) means that for each tumour the width of the proliferating rim decreases until it reaches a steady state and as such C represents a ‘local maximum‚. Figure 12 shows the corresponding heatmaps for both the steady state volumes and volume fractions for each constituent of the tumour spheroid.

## References

Ahmed KA, Correa CR, Dilling TJ, Rao NG, Shridhar R, Trotti AM, Wilder RB, Caudell JJ (2014) Altered Fractionation Schedules in Radiation Treatment: A Review. Seminars in Oncology 41(6):730–750, DOI 10.1053/j.seminoncol.2014.09.012, URL http://linkinghub.elsevier.com/retrieve/pii/S0093775414002322

Alper T, Howard-Flanders P (1956) Role of Oxygen in Modifying the Radiosensitivity of E. Coli B. Nature 178:978–979

Araujo RP, McElwain DLS (2004) A history of the study of solid tumour growth: The contribution of mathematical modelling. Bulletin of Mathematical Biology 66(5):1039–1091, DOI 10.1016/j.bulm.2003.11.002,3843079723

Breward C, Byrne H, Lewis CE (2003) A Multiphase Model Describing Vascular Tumour Growth. Bulletin of Mathematical Biology65(4):609–640, DOI 10.1016/S0092-8240(03)00027-2, URLhttp://link.springer.com/10.1016/S0092-8240(03)00027-2

Byrne H, Chaplain M (1998) Necrosis and apoptosis: distinct cell loss mechanisms in a mathematical model of avascular tumour growth. Journal of Theoretical Medicine 1:223–235, DOI 10.1080/10273669808833021, URL http://www.tandfonline.com/doi/abs/10.1080/10273669808833021

Byrne HM (2012) Continuum models of avascular tumour growth. In: Antoniouk AV, Melnik RVN (eds) Mathematics and Life Sciences, chap 12, pp 279–312

Byrne HM, King JR, McElwain DLS, Preziosi L (2003) A two-phase model of solid tumour growth. Applied Mathematics Letters 16(4):567–573, DOI 10.1016/S0893-9659(03)00038-7

Carlson DJ, Stewart RD, Semenenko VA (2006) Effects of oxygen on intrinsic radiation sensitivity: A test of the relationship between aerobic and hypoxic linear-quadratic (LQ) model parameters. Medical physics 33(9):3105–3115, DOI 10.1118/1.2229427

Enderling H, Chaplain MAJ, Hahnfeldt P (2010) Quantitative Modeling of Tumor Dynamics and Radiotherapy. Acta Biotheoretica 58(4):341–353, DOI 10.1007/s10441-010-9111-z

Eschrich SA, Pramana J, Zhang H, Zhao H, Boulware D, Lee JH, Bloom G, Rocha-Lima C, Kelley S, Calvin DP, Yeatman TJ, Begg AC, Torres-Roca JF (2009) A Gene Expression Model of Intrinsic Tumor Radiosensitivity: Prediction of Response and Prognosis After Chemoradiation. International Journal of Radiation Oncology Biology Physics 75(2):489–496, DOI 10.1016/j.ijrobp.2009.06.014

Folkman J, Hochberg M (1973) Self-regulation of growth in three dimensions. Journal of Experimental Medicine 138(4):745–753, DOI 10.1084/jem.138.4.745 http://www.jem.org/cgi/doi/10.1084/jem.138.4.745

Fowler JF (2006) Development of radiobiology for oncology-a personal view. Physics in medicine and biology 51(13):R263–R286, DOI 10.1088/0031-9155/51/13/R16

Frieboes HB, Lowengrub JS, Wise S, Zheng X, Macklin P, Bearer EL, Cristini V (2007) Computer simulation of glioma growth and morphology. NeuroImage 37:S59–S70, DOI 10.1016/j.neuroimage.2007.03.008, URL http://linkinghub.elsevier.com/retrieve/pii/S1053811907001784

Greenspan H (1972) Models for the Growth of a Solid Tumor by Diffusion

Grimes DR, Kelly C, Bloch K, Partridge M (2014) A method for estimating the oxygen consumption rate in multicellulartumourspheroids. Journalof the Royal Society, Interface 11(92):20131,124, DOI 10.1098/rsif.2013.1124, URL http://rsif.royalsocietypublishing.org/content/11/92/20131124.short

Hendry JH (1979) Radiobiology for the Radiologist, vol 39. DOI 10.1097/00003072-199501000-00029

Hirschhaeuser F, Menne H, Dittfeld C, West J, Mueller-Klieser W, Kunz-Schughart LA (2010) Multicellular tumor spheroids: An underestimated tool is catching up again. Journal of Biotechnology 148(1):3–15, DOI 10.1016/j.jbiotec.2010.01.012, URL http://linkinghub.elsevier.com/retrieve/pii/S0168165610000398

Joiner M, van der Kogel A (2009) Basic Clinical Radiobiology Fourth Edition. CRC Press, DOI 10.1201/b13224

Marcu LG (2010) Altered fractionation in radiotherapy: From radiobiological rationale to therapeutic gain. Cancer TreatmentReviews 36(8):606–614, DOI 10.1016/j.ctrv.2010.04.004, URL http://linkinghub.elsevier.com/retrieve/pii/S0305737210000824

McAneney H, O’Rourke SFC (2007) Investigation of various growth mechanisms of solid tumour growth within the linear-quadratic model for radiotherapy. Physics in medicine and biology 52(4):1039–1054, DOI 10.1088/0031-9155/52/4/012

Mueller-Klieser W (1987) Multicellular spheroids -A review on cellular aggregates in cancer research. Journal of Cancer Research and Clinical Oncology 113(2):101–122, DOI 10.1007/BF00391431

Muriel VP (2006) The impact on oncology of the interaction of radiation therapy and radiobiology. Clinical and Translational Oncology 8(2):83–93, DOI 10.1007/s12094-006-0163-0

Nilsson P, Thames HD, Joiner MC (1990) A generalized formulation of the ‘incomplete-repair’ model for cell survival and tissue response to fractionated low dose-rate irradiation. International journal of radiation biology 57(1):127–142, DOI 10.1080/09553009014550401

O’Rourke SFC, McAneney H, Hillen T (2009) Linear quadratic and tumour control probability modelling in external beam radiotherapy. Journal of Mathematical Biology 58(4-5):799–817, DOI 10.1007/s00285-008-0222-y

Preziosi L, Tosin A (2009) Multiphase modelling of tumour growth and extracellular matrix interaction: Mathematicaltoolsandapplications. Journal of Mathematical Biology 58(4-5):625–656, DOI 10.1007/s00285-008-0218-7

Prokopiou S, Moros EG, Poleszczuk J, Caudell J, Torres-Roca JF, Latifi K, Lee JK, Myerson R, Harrison LB, Enderling H (2015) A proliferation saturation index to predict radiation response and personalize radiotherapy fractionation. Radiation Oncology 10(1):159, DOI 10.1186/s13014-015-0465-x, URL http://www.ro-journal.com/content/10/1/159

Sachs RK, Hahnfeld P, Brenner DJ (1997) The link between low-LET dose-response relations and the underlying kinetics of damage production/repair/misrepair. International journal of radiation biology 72(4):351–374, DOI 10.1080/095530097143149

Sachs RK, Hlatky LR, Hahnfeldt P (2001) Simple ODE models of tumor growth and anti-angiogenic or radiation treatment. Mathematical and Computer Modelling 33(12-13):1297–1305, DOI 10.1016/S0895-7177(00)00316-2

Schaller G, Meyer-Hermann M (2006) Continuum versus discrete model: a comparison for multicellular tumour spheroids. Philosophical Transactions of the Royal Society A: Mathematical, Physical and Engineering Sciences 364(1843):1443–1464, DOI 10.1098/rsta.2006.1780, URL http://rsta.royalsocietypublishing.org/cgi/doi/10.1098/rsta.2006.1780

Sutherland R, Carlsson J, Durand R, Yuhas J (1981) Spheroids in Cancer Research. Cancer Research 41(July):2980–2984, URL http://cancerres.aacrjournals.org/content/41/7/2980.abstract

Thrall DE (1997) Biologic basis of radiation therapy. The Veterinary clinics of North America Small animal practice 27(1):21–35

Torre LA, Bray F, Siegel RL, Ferlay J, Lortet-tieulent J, Jemal A (2015) Global Cancer Statistics, 2012. CA: a cancer journal of clinicians 65(2):87–108, DOI 10.3322/caac.21262., URL http://onlinelibrary.wiley.com/doi/10.3322/caac.21262/abstract

Withers HR (1999) Radiation biology and treatment options in radiation oncology. Cancer research 59(7 Suppl):1676s–1684s

